# A mutation in monoamine oxidase (MAO) affects the evolution of stress behavior in the blind cavefish *Astyanax mexicanus*

**DOI:** 10.1101/2020.04.09.033266

**Authors:** Constance Pierre, Naomie Pradère, Cynthia Froc, Patricia Ornelas-García, Jacques Callebert, Sylvie Rétaux

## Abstract

The neurotransmitter serotonin controls a great variety of physiological and behavioral processes. In humans, mutations affecting the monoamine oxidase or MAO, the serotonin-degrading enzyme, are highly deleterious. Yet, blind cavefish of the species *A. mexicanus* carry a partial loss-of-function mutation in MAO (P106L) and seem to thrive in their subterranean environment. Here, we established 4 fish lines, corresponding to the blind cave-dwelling and the sighted river-dwelling morphs of this species, with or without the mutation, in order to decipher the exact contribution of *mao* P106L in the evolution of cavefish neuro-behavioral traits. Unexpectedly, although *mao* P106L appeared as an excellent candidate for the genetic determinism of the loss of aggressive and schooling behaviors in cavefish, we demonstrated that it was not the case. Similarly, the anatomical variations in monoaminergic systems observed between cavefish and surface fish brains were independent from *mao* P106L, and rather due other, morph-dependent developmental processes. On the other hand, we found that *mao* P106L strongly affected anxiety-like behaviors. Cortisol measurements showed lower basal levels and an increased amplitude of stress response after a change of environment in fish carrying the mutation. Finally, we studied the distribution of the P106L *mao* allele in wild populations of cave and river *A. mexicanus*, and discovered that the mutant allele was present - and sometimes fixed - in all populations inhabiting caves of the Sierra de El Abra. The possibility that this partial loss-of-function *mao* allele evolves under a selective or a genetic drift regime in the particular cave environment is discussed.

## Introduction

Monoaminergic systems control a variety of vital physiological functions in vertebrates, ranging from stress response (Dinan, 1996; Winberg et al., 1997) to gut motility (Bülbring and Crema, 1958; Gershon, 2013; Mawe and Hoffman, 2013), metabolic homeostasis (El-Merahbi et al., 2015), immune function (Khan and Deschaux, 1997; Nicole and Randy, 2013; Shajib and Khan, 2015) or reproduction (Prasad et al., 2015). They also play roles in brain, heart, ocular and craniofacial development (Baker and Quay, 1969; Moiseiwitsch, 2000; Ori et al., 2013; Sodhi and Sanders-Bush, 2004; Souza and Tropepe, 2011). Moreover and crucially, due to their central neuromodulatory functions, monoaminergic systems control various aspects of animal behavior: aggressiveness (Edwards and Kravitz, 1997; Nelson and Trainor, 2007; Olivier, 2004; Popova, 2006), locomotion (Beninger, 1983; Brocco et al., 2002; Gabriel et al., 2009; Pearlstein, 2013; Perrier and Cotel, 2015), sleep and arousal (Jouvet, 1999; Oikonomou et al., 2019; Scammell et al., 2017), food intake (Pérez-Maceira et al., 2016; Voigt and Fink, 2015), or olfaction (Gaudry, 2018), among others.

Plastic changes in the monoaminergic systems are widely described, and can occur in virtually all animal species during acclimation to variations in the environment such as temperature, altitude, or season (e.g., Stefano and Catapane, 1977; Hernádi et al., 2008; Nakagawa et al., 2016; Vaccari et al., 1978). Evolutionary changes in monoaminergic systems during adaptation to a novel environment are much less studied. However, inter-species and intra-species genetic variations in monoaminergic pathway genes exhibiting different behaviors have been reported (Bergey et al., 2016; Staes et al., 2019) examples in non-human primates). This opens the possibility that genetically encoded, evolutionary changes in monoaminergic pathways could be selected during adaptation of species to their environment.

The fish *Astyanax mexicanus* is an excellent model to study adaptation after an environmental shift. It comes in two inter-fertile forms: sighted and pigmented morphs, which live in rivers of Northern Mexico; and blind depigmented morphs, which live in 31 caves in North-East Mexico (Elliott, 2018; Mitchell et al., 1977). Surface-like, common ancestors of the two extant morphs colonized the caves about 20.000 years ago, and have since then adapted to the total and permanent darkness of the subterranean environment (Fumey et al., 2018; Policarpo et al., 2020).

Surface fish (SF) and cavefish (CF) display many morphological, behavioral and physiological differences. The eyes of CF rapidly degenerate during development (Hinaux et al., 2015; Jeffery, 2009), and their brains show multiple differences when compared to river-dwelling conspecifics (Rétaux et al., 2016). They also have more teeth and modifications of their craniofacial bones structure (Atukorala et al., 2013; Gross et al., 2014; Yamamoto et al., 2003). CF are albino as they do not synthetize melanin (Bilandžija et al., 2013; McCauley et al., 2004), they have a low metabolic rate (Hüppop, 1986; Moran et al., 2014; Salin et al., 2010), and an altered intestinal motility (Riddle et al., 2018). Behavioral differences are manifold: most CF are non-aggressive (Elipot et al., 2013; Espinasa et al., 2005), they do not school (Kowalko et al., 2013), they sleep very little (Duboué et al., 2011), and they show intense exploratory behavior (Patton et al., 2010). They are attracted to vibrations (Yoshizawa et al., 2010) and have an exceptional sense of smell (Hinaux et al., 2016). This ensemble is called the “cavefish behavioral syndrome” and, like most anatomical and physiological modifications cited above, is often considered as adaptive for cave life.

At the molecular level, cavefish originating from the Pachón cave carry a coding point mutation in the gene for *Monoamine Oxidase* (*mao*; P106L mutation), the serotonin degrading enzyme (Elipot et al., 2014). P106L decreases MAO enzymatic activity, which results in enhanced monoamine levels in the CF brain – note that fish possess only one form of MAO, contrarily to mammals. Because of the many roles of monoaminergic systems in development, physiology and behavior, the P106L *mao* mutation could be responsible for some of the CF special traits and could participate in their adaptation to the cave environment.

Here, we have systematically analyzed the phenotypic effects of the P106L *mao* mutation at the neuroanatomical, behavioral and physiological levels. We have also studied the distribution and the evolutionary history of the P106L mutation, in order to discuss its adaptive nature and its impact on cavefish evolution.

## Methods

### Fish husbandry

Laboratory stocks of *A. mexicanus* surface fish (origin: Texas) and cavefish (Pachón population) were obtained in 2004–2006 from the Jeffery laboratory at the University of Maryland, College Park, MD, USA and were since then bred in our local facility (Gif sur Yvette, France). Fishes were maintained at 23–26°C on 12:12-h light: dark cycle and they were fed twice a day with dry food. The fry were raised in Petri dishes and fed with micro-worms after opening of the mouth. Animals were treated according to the French and European regulations for handling of animals in research. SR’s authorization for use of *Astyanax mexicanus* in research is 91-116 and the Paris Centre-Sud Ethic Committee protocol authorization number related to this work is 2017-05#8604. The animal facility of the Institute received authorization 91272105 from the Veterinary Services of Essonne, France, in 2015.

### Wild fish samples

our fin clips collection from wild cavefish and surface fish populations were obtained during field sampling campaigns between 2013 and 2019, under the auspices of the field permits 02241/13, 02438/16, 05389/17 and 1893/19 delivered by the Mexican Secretaría del Medio Ambiente y Recursos Naturales to POG and SR.

### Fish lines

To obtain the Pachón CF without the P106L mutation, we crossed heterozygotes fish identified among our lab Pachón breeding colony (Fig. 1A), taking advantage that the P106L mutation is not fixed in the Pachón population. To obtain a SF line carrying the mutation, a cross between a SF (wild type *mao*) and a CF carrying the P106L (homozygote mutant) was followed by 4 backcrosses with SF (Fig. 1B). At each generation, we selected fish carrying the P106L mutation (heterozygotes) and backcrossed them with SF to dilute the cave genome little by little. Then, to obtain SF homozygote mutants, we intercrossed the last generation together. Theoretically, after 5 generations, the percentage of cave genome is 3.13%. Note that the generation time between the spawn of the n generation and the spawn of the n+1 generation was about 8 months.

**Figure 1.**
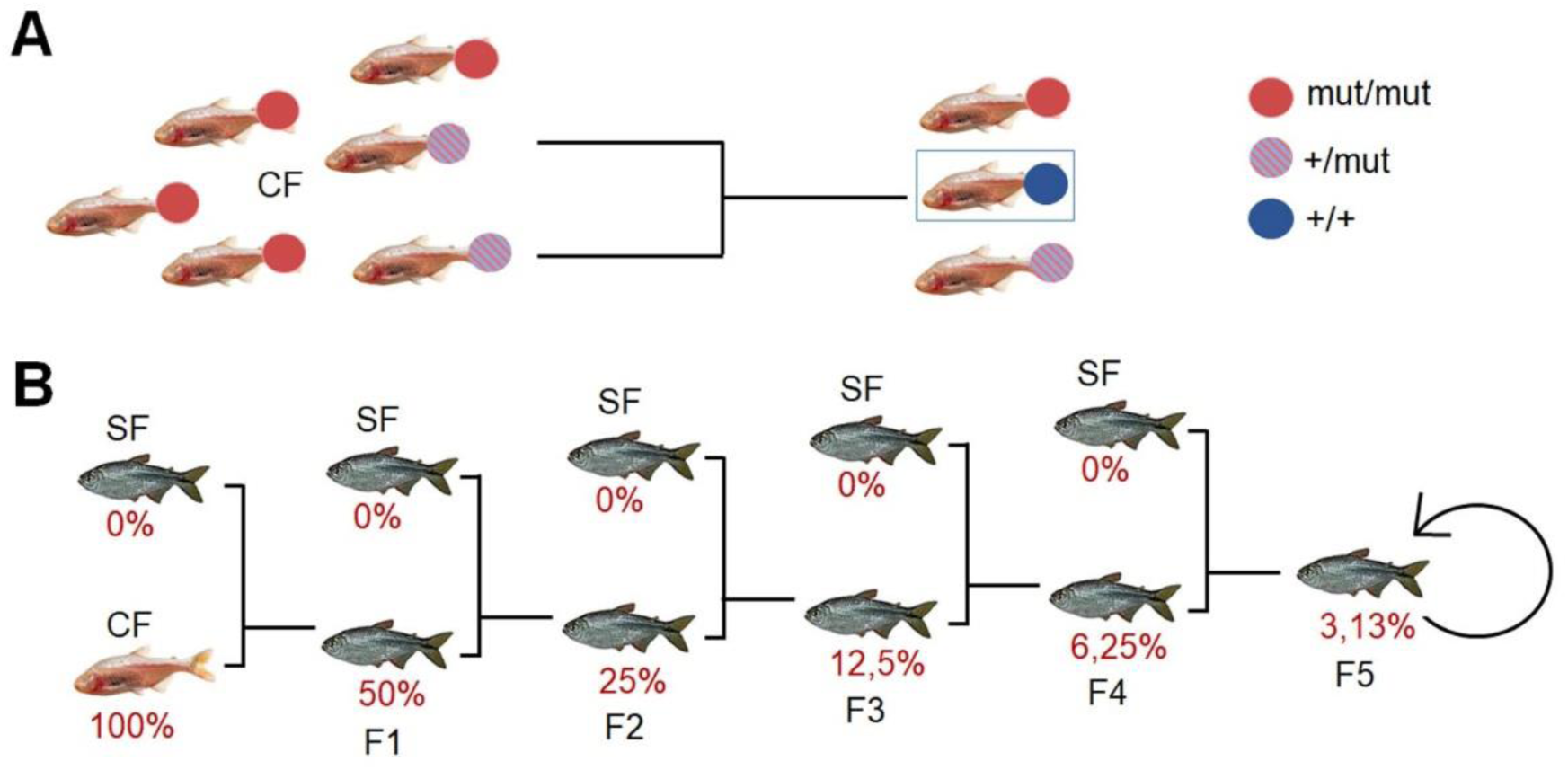
Generation of the lines of interest by crosses. (A) Generation of cavefish (CF) without the P106L *mao* mutation by crosses of heterozygotes. The dots on fishes indicate their genotype: mutant (mut/mut; red), non-mutant (+/+; blue) or heterozygote (+/mut; striped). (B) Generation of surface fish (SF) with the P106L *mao* mutation by backcrosses. The percentage in red means the theoretical percentage of cavefish genome at each generation.

### *mao* P106L allele genotyping

wild fin-clips preserved in ethanol 100% and lab fin clips were genotyped as follows. We performed a crude lysis with proteinase K in lysis buffer (100mM Tris; 2mM EDTA; 0.2% Triton; 0.01µg/µl PK), followed by PCR (primer F-GGGAAATCATATCCATTCAAGGGG; primer R-CTCCATGCCCATCTTGTCCATAG) and a purification of DNA (NucleoSpin® Gel and PCR Clean-up). We used the sequencing service of Eurofins Genomics. Homozygotes and heterozygotes at position 106 were easily detected and identified on sequence chromatograms (see Fig. 2B).

**Figure 2.**
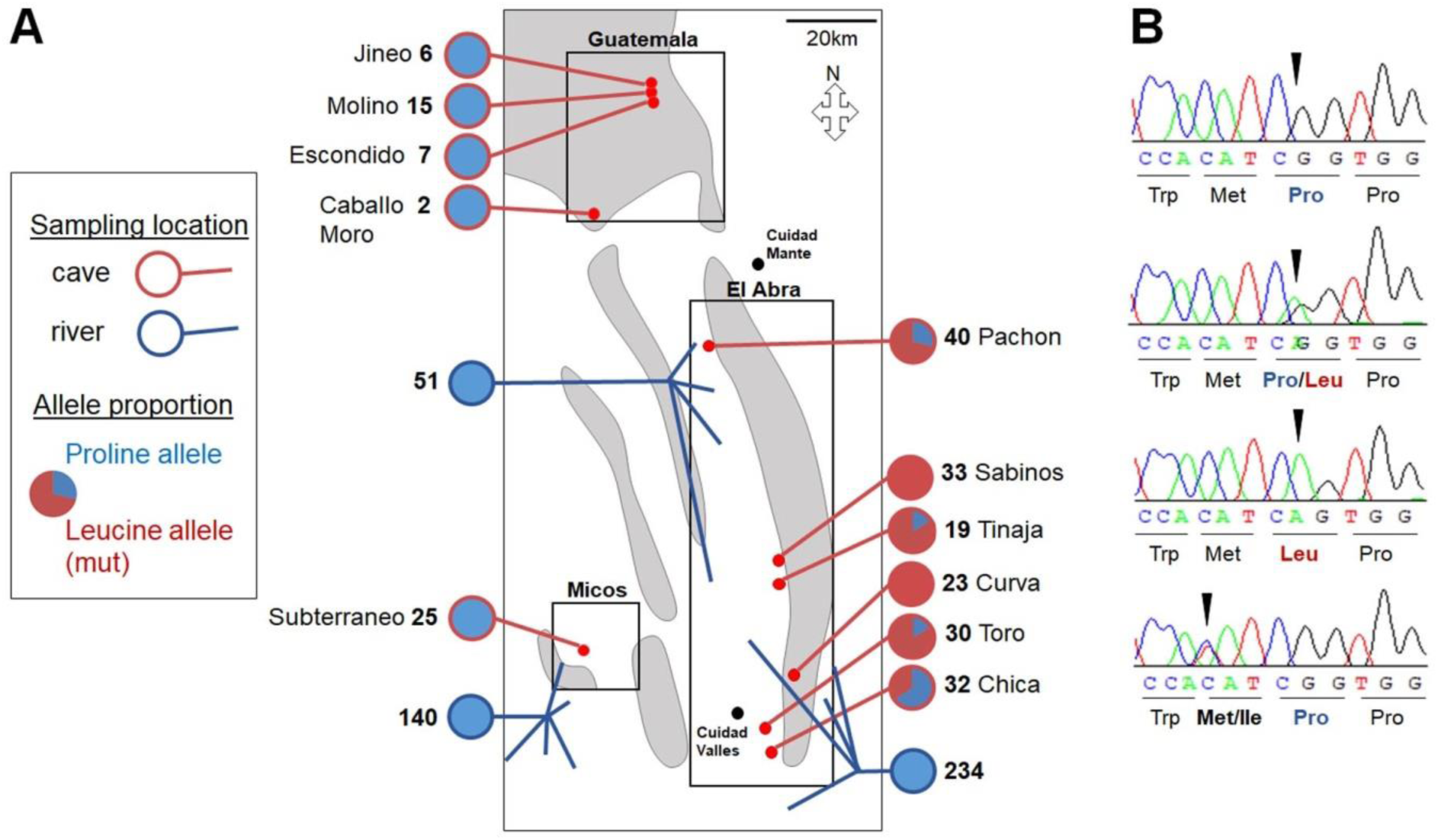
Distribution of *mao* alleles in the wild. (A) The proportion of mutant alleles is given for 11 caves (red edges) and 3 zones of sampling in surface including (total 13 locations, blue edges). The number of fish sampled is indicated for each location. (B) Sequence chromatograms centered on MAO amino acid 106. From top to bottom, a non-mutant individual, a heterozygote, a mutant homozygote (P106L), and last, a surface fish with heterozygous M107I mutation. The reverse sequence is shown.

### *mao* genomic fragment sequencing

a fragment spanning 3838bp on the *mao* gene was sequenced in 110 wild-sampled individuals originating from different caves and surface rivers. The fragment includes the exons n°4 (where the P106L mutation is located), 5, and 6, and is comprised between the positions NC_035915.1:32708790 and NC_035915.1:32713975. This fragment was subdivided in 4 smaller fragments, which were easier to sequence. The primers used were: Fw1-TTGTGCCTCTGTGGTGATGA, Rv1-AGTGCCGGAACCTAAAGGA; Fw2-AGCTCGCTAGCTAAATGTGTGA,Rv2-GAAGAATGCTTGCTGGAGCTG; Fw3-TCTCATCTGCTTGTTGATGGCT, Rv3-TCCCCTAGGAGCAACGAAAC; Fw4-TTTTATGGTGGCATGCAGAAGTG, Rv4-ATGATACTGCAAGCGAAGCC. We used the sequencing service of Eurofins Genomics.

### Immunofluorescence, immunohistochemistry and imaging

6dpf larvae were fixed in 4% paraformaldehyde in phosphate-buffered saline, gradually deshydrated in ethanol, and permeabilized 30min in EtOH: Xylen 1:1 at RT, then 10min in EtOH; aceton 1:2 at -20°C. The samples were rehydrated in phosphate-buffered saline and 1% tween (PBST). In addition to that, for immunohistochemistry, samples were incubated in PBST with 6% H2O2, 30min at RT. Next, the inferior jaw of the larvae were removed to make the ventral brain more accessible to antibodies. The samples were incubated in blocking buffer (10% sheep serum, 1% triton, 1% tween 20, 1% DMSO, 1% BSA in PBST) for 2h, then incubated with the primary antibody (rat anti-serotonin, Chemicon MAB352, 5/1000; rabbit anti-tyrosine hydroxylase, Sigma Aldrich SAB2701683, 2/1000) in buffer (1% sheep serum, 1%triton, 1%DMSO, 1%BSA in PBST) for 3 days, followed by the secondary antibodies (biotinylated goat anti-rat antibody, Jackson 112-066-072, 1/500; goat anti-rabbit Alexa fluor 635, Life technologies A31577, 1/500) and DAPI (4/1000) for immunofluorescence. For immunohistochemistry, revelation was performed using an avidin/biotin complex coupled with peroxidase (ABC kit, Universal Vectastain Kit PK6200) and diaminobenzidine (Sigma Aldrich).

Brains were dissected by removing the palate and mounted in Vectashield (Vector Laboratories) and glycerol for immunofluorescence and immunohistochemistry, respectively. Images were taken on a Leica confocal Sp8 and a macrozoom Nikon AZ100 for immunofluorescence and immunohistochemistry, respectively. Images were corrected for brightness and contrast in ImageJ or Photoshop but no other corrections were made.

### Behavioral analyses

#### Aggressiveness

3 months-old fish were used in a resident-intruder test. Fish were placed individually in tanks (12×9cm) with 200ml of water for the night. In the morning, intruders were transferred in tanks of resident, and a video of 1h was recorded. Attacks were counted manually using the software ODREC (Observational Data Recorder).

#### Shoaling

Groups of 6 fish were put in tanks of 25×18cm (1.2l) for 2months-old fishes, and 40×23cm (3.6l) for 5months-old fishes. Tanks were placed on an infrared floor. A video of 10min was recorded after 10min of habituation, and the inter-individual distances (IID) and nearest-neighbor distances (NND) were calculated with the ViewPoint software. IID and NND were simulated for a random distribution of the fish with a program writer under Scilab 5.5.2. Of note, here we used the term “shoaling” (as opposed to “schooling”) because we did not measure angles between fish.

#### Locomotion

6dpf larvae were placed individually in 24-wells plates, 2 months-old fish in 12×9cm tanks (200ml), and 5 months-old in 19×10cm (600ml). Tanks were placed on an infrared floor. Videos were recorded after 10, 30min, 1h or 24h of habituation, depending on the experiment. For the measure of locomotion in groups, groups of six 5months-old fish were placed in tanks of 40×23cm (5l) on infrared floors, with a 12:12h light:dark cycle. Videos of 10min were recorded.

#### Food intake

1 year-old fish were placed individually in tanks (25×11×10cm) for 5 days with no food available. The 4^th^ day, they were weighted. The 5^th^ day, food was added in tanks, and the fish were weighted 1h after feeding. The same protocol was used to measure food intake in groups, except that the fish were in groups of 20, and in their home tank at the fish facility.

#### Anxiety behaviors

the same protocol as for the recording of locomotion was used for 5 month-old fish, alone in their tank. The different stress behaviors were analyzed manually with ODREC. Freezing: the fish is immobile, paralyzed, and sometimes loses equilibrium and stays on the side. Thigmotaxis: fast swimming, head against the wall of the tank. Erratic movements: the fish swims very fast, and frequently changes of direction, with angles of 90°. Attempts to dive: the fish swims vertically with frenzy, head against the bottom of the tank, sometimes moving horizontally along the tank, sometimes not. Freezing, thigmotaxis and erratic movements are described and widely used to measure anxiety in other fish (Blaser and Gerlai, 2006; Blaser et al., 2010; Cachat et al., 2010; Champagne et al., 2010; Maximino et al., 2010; Schnörr et al., 2012). Although bottom dwelling was described as an anxiety behavior (Cachat et al., 2010; Chin et al., 2018; Egan et al., 2009; Levin et al., 2007), to our knowledge attempts to dive were not, and could actually correspond to a form of thigmotaxis as they represent efforts to escape by the bottom of the tank.

### Cortisol and monoamine levels

6dpf larvae were anesthetized in water at 1°C. The heads were immediately cut and recovered to measure noradrenalin, and the bodies were recovered to measure noradrenalin and adrenalin. A sample (n=1) was formed with 15 heads or 15 bodies, in 400µl of HCl (10^−3^M). Adults were placed by 4-6 in tanks (fish facility home tanks for several weeks, novel tank for 10min or 24h). They were quickly anesthetized in water at 1°C, and immediately frozen to avoid losing blood during dissection. Then, the head was cut and the brain dissected out and placed in 400µl of HCl (10-3M) to measure monoamines, while the body (with gills) was placed in 1ml of HCl (10^−3^M) to measure cortisol. Before analysis, tissues were crushed and centrifuged at 20.000g for 1h. The supernatant was analyzed by fluorimetry for serotonin (HPLC), coulometry for catecholamines (HPLC), and immune-chemo-luminescence for cortisol (Abott Alinity).

### Cardiac rhythm

8dpf larvae were anesthetized in a 140ppm tricaine solution, and the cardiac beats were counted manually. We cannot rule out that tricaine had a bradycardic effect. However, if any, the effect was the same on all fish lines. Of note, the concentration of tricaine used here was low and this method was preferred to, e.g., low temperature, which had a clear and immediate effect on cardiac frequency.

### Statistical analyses

No statistical method was used to predetermine sample size. No data were excluded from analysis, and sample allocation was random after genotyping. Sex was not considered in the analyses as it impossible to determine sex in *Astyanax mexicanus* before the age of 6-7 months without dissection/sacrifice. Analyses (except automated analyses by Viewpoint software) could not be blinded: for anatomical analyses, brains from SF or CF are easily recognizable by eye size, and for behavioral analyses, the investigator can also easily recognize the morphs on the videos. All experiments (except anatomy for mutant SF and their siblings in Fig. 3 and feeding assay in group in Fig. 6) were reproduced at least twice (=technical replicates), and the results were pooled (=biological replicates, corresponds to the n values indicated on graphs). Statistical analyses were performed with BiostaTGV (https://biostatgv.sentiweb.fr/), using non-parametric Mann-Whitney tests (normal distribution was not tested). All graphs show mean±sem, and the number of samples included is systematically indicated on graph bars. P values are shown. In all figures, *** p<0.0001, ** p<0.001 and * p<0.01.

**Figure 3.**
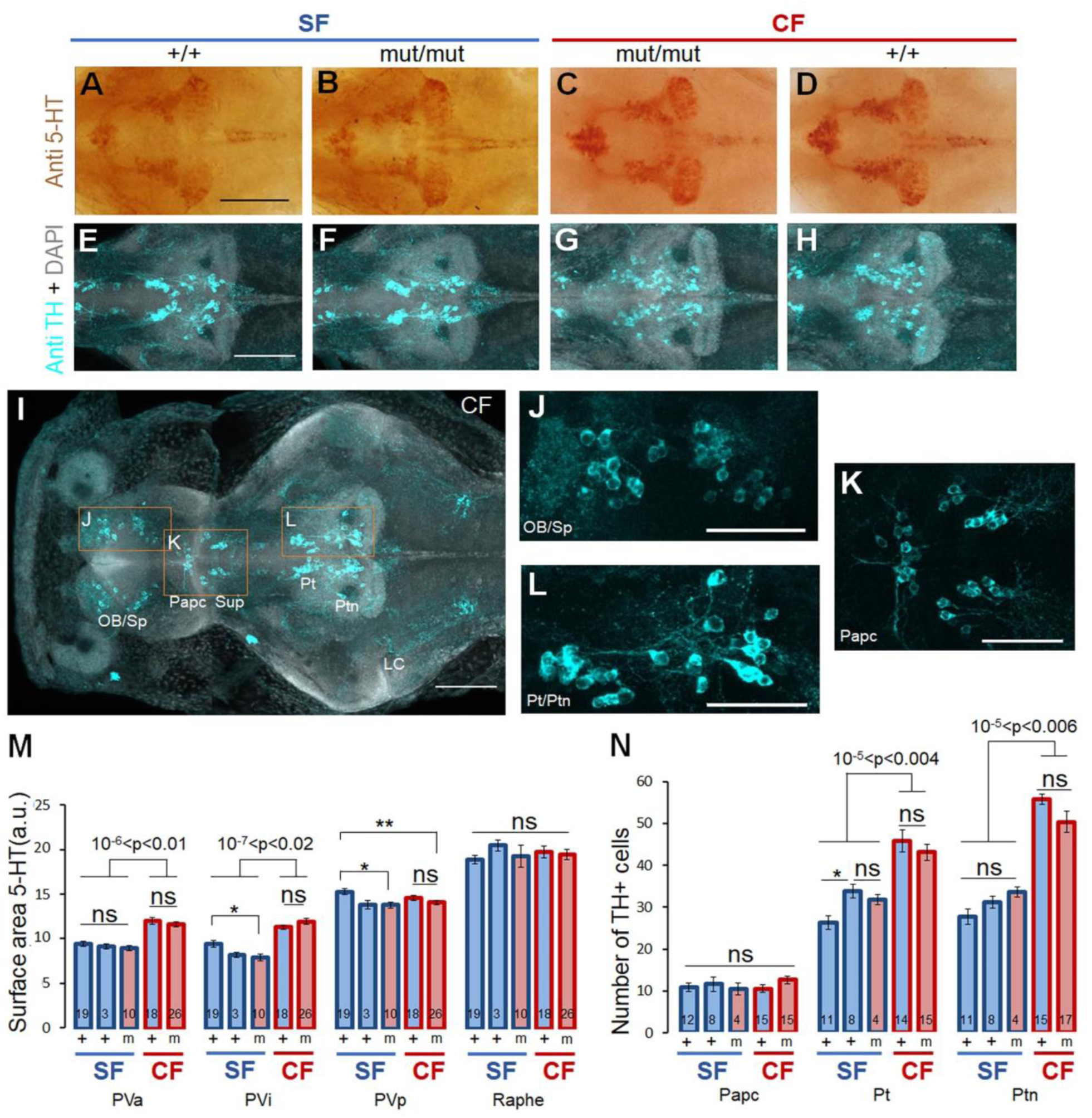
5-HT and DA systems in 6dpf larvae. (A-H) Immunohistochemistry for 5-HT (A-D) and immunofluorescence for TH (E-H) in brains of the 4 lines. Scale: 100µm. (I) Image of a CF whole-brain, showing a general view of TH-positive cells. Scale: 100µm. (J-L) High magnification views of TH-positive clusters of olfactory bulb and subpallium (OB/Sp) (J), Preoptic anterior parvo-cellular (Papc) and suprachiasmatic (Sup) clusters (K), and posterior tuberculum (Pt) and posterior tuberal nucleus (Ptn) (L). Scale: 50µm. LC: Locus coeruleus. (M) Quantification of 5-HT groups areas, for the indicated genotypes. PVa/i/p stands for anterior/intermediate/posterior paraventricular nucleus. (N) Number of TH-positive cells in Papc, Pt and Ptn. Non-mutant SF (+) shown are either wildtype (first bar of each group), or mutants siblings (second bar).

**Figure 4.**
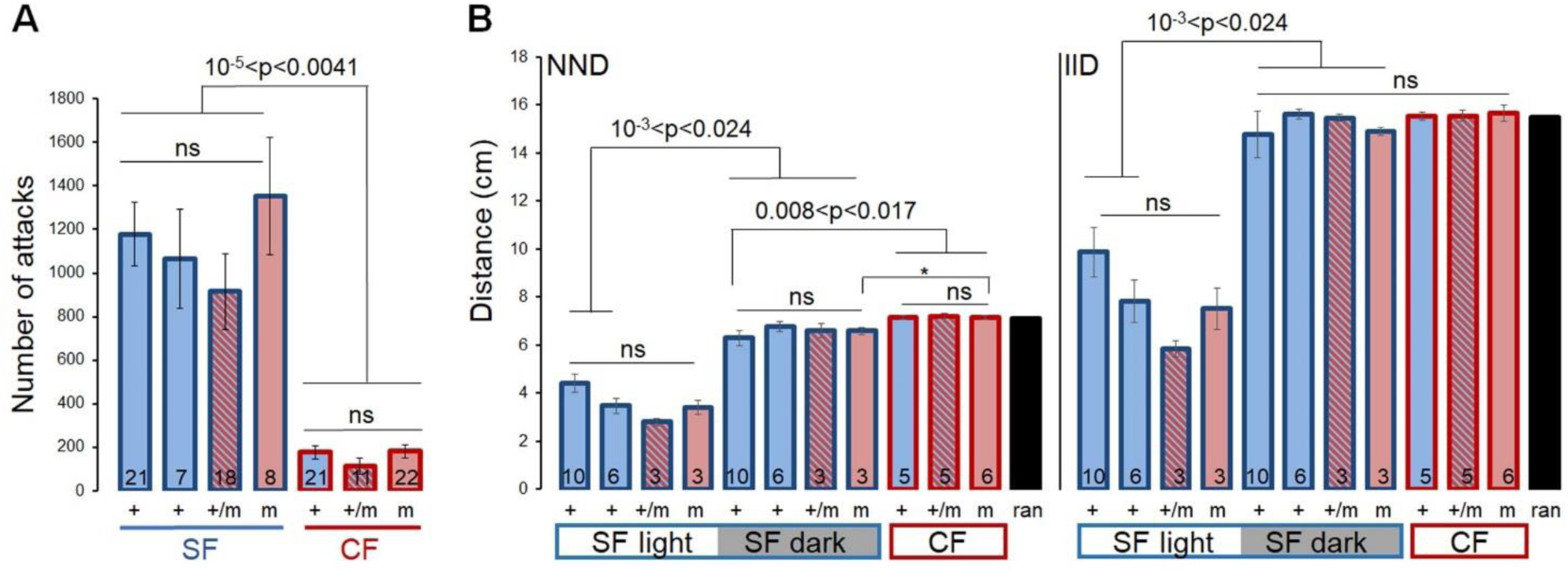
Aggression and schooling. (A) Number of attacks counted during 1h for SF and CF of the indicated genotypes. Non-mutant SF (+) tested are either wildtype (first bar of each group), or mutants siblings (second bar). (B) Nearest-neighbor distance (NND) and inter-individual distances (IID) in groups of fish, measured in the light (SF and CF) or in the dark for SF. N=1 corresponds to a group of 6 fish of the same morph and genotype. NND and IID were calculated from 10min recordings. The values of NND and IID for a random distribution of fish (ran) was calculated with 100.000 positioning simulations of 6 points (=fishes) in a square of the dimensions of the real tank.

**Figure 5.**
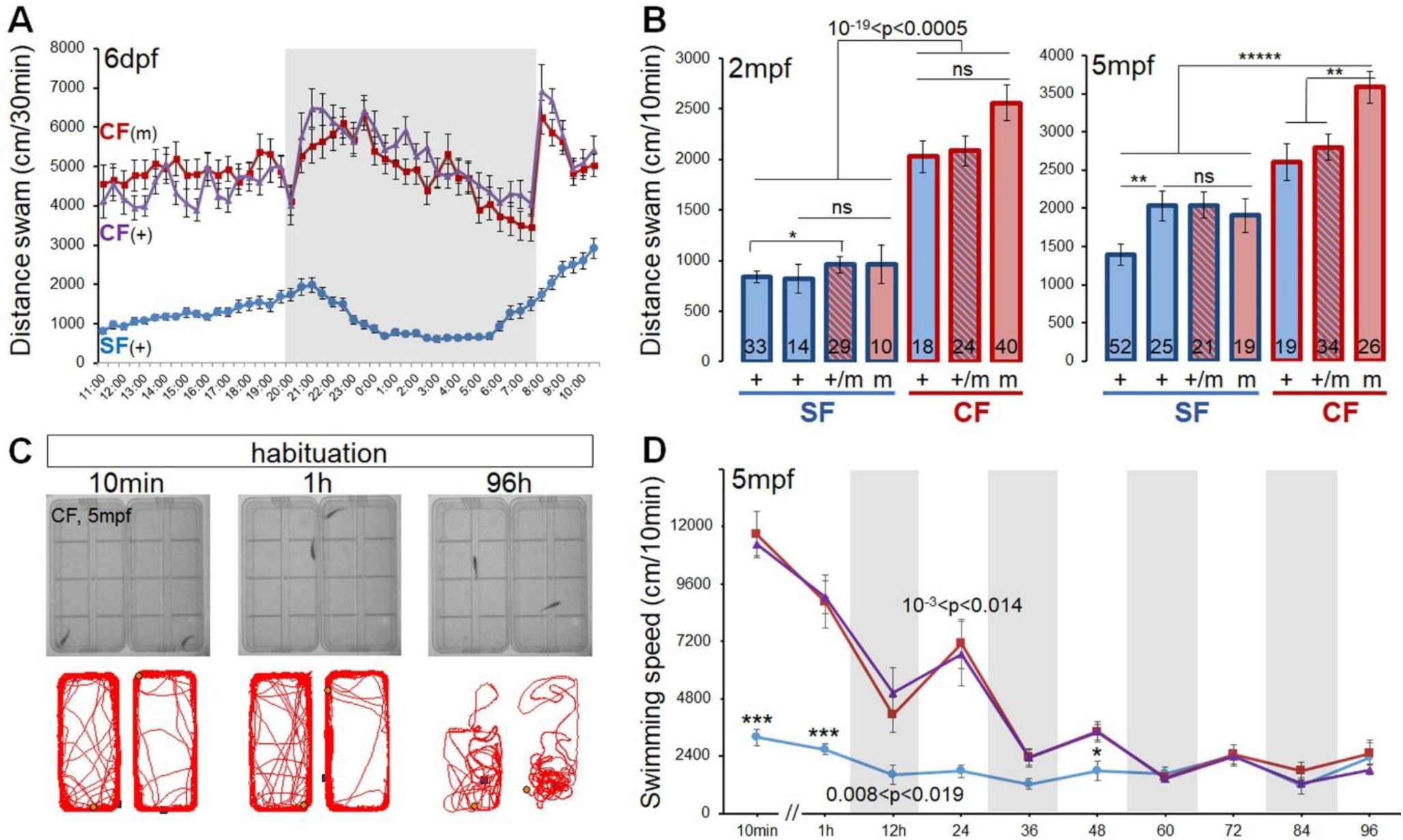
Locomotion. (A) Distance travelled by 6dpf larvae in SF (blue; n=46), P106L homozygotes mutants CF (red; n=30) and non-mutant CF (purple; n=26). The recordings lasted 24h, with a 8AM:8PM day:light cycle. (B) Distance travelled by 2 or 5 month-old fish in 10min, 1h after the beginning of the experiment. The fish lines tested are indicated. Non-mutant SF (+) tested are either wildtype (first bar), or mutants siblings (second bar). (C) Images of 5 month-old CF in the test tank and their trajectory during 10min periods, at 10min, 1h and 96h after the beginning of the experiment. (D) Absolute speed measured during 10min recordings for groups of six 5 month-old fish. Recordings were performed on the day and the night during 5 days. Fishes tested are SF (blue; n=5), P106L mutant homozygotes CF (red; n=10), and non-mutant CF (purple; n=12).

**Figure 6.**
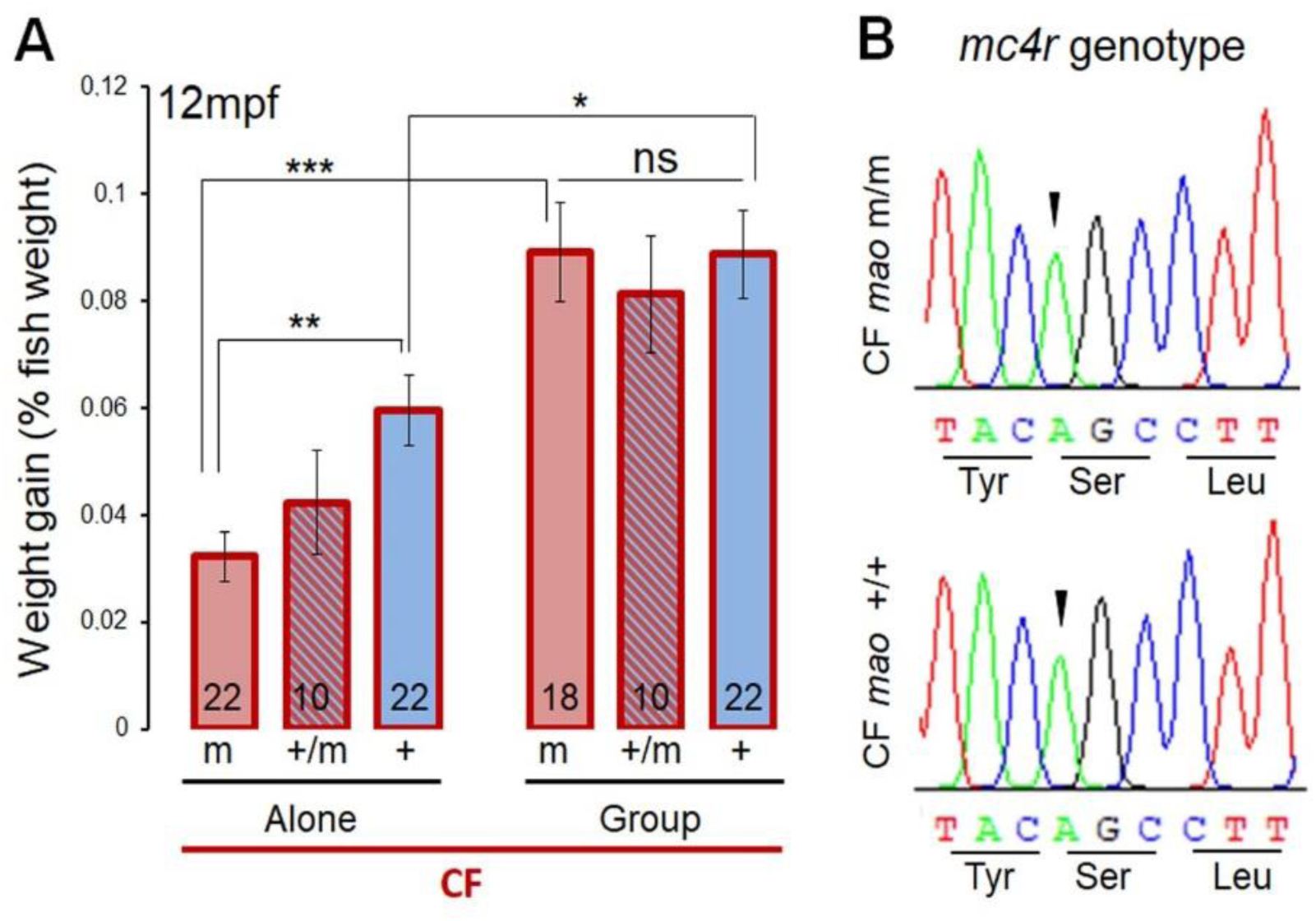
Food intake in CF and *mc4r* genotype. (A) Weight gain in 1-year old CF, either alone with 5 days of habituation, or in groups in their home tank at the fish facility. The same fish (genotypes indicated) were tested in the 2 conditions. (B) Sequence chromatograms for *mc4r* centered on the Gly-to-Ser mutation of a P106L *mao* mutant homozygote CF (top), and a non-mutant CF (bottom). The two fish carry the homozygous *mc4r* mutation.

## Results

### Distribution of *mao* alleles in wild *A. mexicanus* cavefish populations

To estimate the spread of *mao* P106L in natural populations, and its potential relevance to cave adaptation, we analyzed fin clips sampled during field expeditions. We genotyped 232 cavefish from 11 different caves and 425 surface fish from several rivers and water points in the region of Ciudad Valles, SLP, México (Fig. 2; exact locations available on request). The caves can be subdivided into 3 geographical groups: Micos, Guatemala and El Abra, in which the different CF populations were hypothesized to derive from independent colonization events (Bradic et al., 2012). The P106L *mao* mutant allele was present in all sampled caves of the El Abra group (from North to South: Pachón, Tinaja, Los Sabinos, Curva, Toro and Chica), but not in caves of the Micos and Guatemala groups (Fig. 2A). Whereas the wildtype allele was recovered in Pachón, Tinaja, Toro and Chica CF populations, all fish sampled in Curva (n=23) and Los Sabinos (n=33) were homozygous for the mutant allele. This suggested that the mutation has reached fixation in theses 2 caves. The mutant *mao* allele was found in high proportions in all El Abra caves except Chica (1/3 of mutant alleles).

On the other hand, the P106L mutation was never found in the 425 genotyped SF (=850 alleles). Of note, in the *mao* exon 4 which contains the P106L mutation and in exons 5/6 which were also amplified for additional genotyping, we identified another synonymous mutation (not shown) as well as a non-synonymous mutation at position 107 (M107I, adjacent to P106L) (Fig. 2B). This indicated the presence of polymorphisms in *mao* in the natural river populations of *A. mexicanus* (see also Suppl Fig. 6 and Discussion). Thus, the P106L mutation is either absent from the SF populations, or present at a frequency lower than 1/850.

To study the phenotypic effects of the P106L *mao* mutation, below we systematically compared 4 lines of fish: SF and Pachón CF, wild type or mutant/P106L. To generate these lines, we took advantage of the presence of heterozygous individuals in our Pachón CF laboratory colony: Pachón CF without the mutation were obtained by crosses between 2 heterozygotes (Fig. 1A), and SF carrying the mutation were obtained by successive backcrosses (Fig. 1B and see methods).

### *mao* P106L does not affect the neuroanatomy of monoaminergic systems

The P106L mutation significantly decreases MAO enzymatic activity, resulting in an increase of monoamine levels in brain and body (Elipot et al., 2014; Pierre et al., in prep). As serotonin can modulate neurogenesis, neuronal differentiation, and synaptogenesis of the serotonergic system and its targets (Lauder, 1993; Pérez et al., 2013; Urtikova et al., 2009; Vitalis et al., 2013; Whitaker-azmitia et al., 1996), and as anatomical variations are reported in the serotonergic system between the two *A. mexicanus* morphs (Elipot et al., 2013), we asked if the P106L *mao* mutation could be responsible for these neuroanatomical changes. We also analyzed the distribution of dopaminergic and noradrenergic neurons, because links between the different monoaminergic systems have been reported (Di Giovanni et al., 2008; Di Giovanni et al., 2010). Serotonin and catecholamine neurons were labelled using immunohistochemistry against 5-HT (serotonin) and TH (tyrosine hydroxylase), respectively, on 6dpf larvae of the 4 different lines.

The serotonergic system in fish is composed of 3 clusters of neurons in the hypothalamus plus the rhombencephalic raphe. In agreement with our previous studies (Elipot et al., 2013), the size of the PVa (anterior paraventricular nucleus) was larger in CF than in SF (Fig. 3A-D, M). Here, we found that the PVi (intermediate nucleus) was also larger in CF. However, the sizes of these 5-HT clusters were similar in P106L mutants (SF or CF) and their non-mutant counterparts. This showed that the P106L mutation does not modify the size of the clusters (a proxy of neuron numbers), and further suggested that the observed difference in PVa and PVi sizes between SF and CF are rather morph-dependent. The fish dopaminergic system is more diffuse than the serotonergic system, and is organized in many discrete clusters showing different cell shapes and labelling intensities (Rink and Wullimann, 2002) (Fig. 3E-K). We counted cell numbers in some of them. For some clusters like the anterior preoptic parvocellular group, the number of cells was the same for all 4 studied lines. For some others like the posterior tuberal nucleus, CF possessed more cells than SF (Fig. 3N). However, similarly to the 5-HT system, we could not detect any influence of the P106L mutation on the number of cells or the organization of the dopaminergic and noradrenergic systems (Fig. 3N, see also Fig. Supp. 1).

In sum, these data suggested that the differences observed between SF and Pachón CF in serotonin PVa and PVi sizes, or else in posterior tuberculum and telencephalic dopaminergic cell numbers are not due to P106L in *mao* but to another, morph-dependent developmental factor.

### The *mao* P106L mutation and the cavefish “behavioral syndrome”

A major difficulty inherent to studies comparing behaviors in the two morphs of *Astyanax* is to disentangle the contribution of blindness and genetics for the expression and evolution of behavioral traits in CF. Indeed, numerous behaviors are visually-driven. Below, using the powerful approach allowed by the comparison of the effects of the P106L mutation in the 4 generated lines, we could bring definitive answers to some long-standing questions.

Previous studies had shown that deprenyl, a selective MAO inhibitor, decreases aggressive and schooling behaviors in SF (Elipot et al., 2013; Kowalko et al., 2013). As these behaviors are typically lost in CF and are part of its so-called “behavioral syndrome”, we first tested the possibility that the P106L *mao* mutation could be responsible for their evolution in CF.

Aggressiveness was assessed using a resident-intruder test, by counting the number of attacks performed between 2 individuals in one hour. As previously described, SF were way more aggressive than Pachón CF (Fig. 4A) (Elipot et al., 2013). However, there was no difference in aggressiveness between mutated and non-mutated SF, nor between mutated or non-mutated CF. We concluded that the P106L *mao* mutation has no influence on the evolution of aggressive behavior in *A. mexicanus*.

Similarly, we analyzed shoaling by measuring inter-individual distance (IID) and nearest neighbor distance (NND) in groups of 6 fish (Fig. 4B). Consistent with previous reports (Kowalko et al., 2013), in SF IID and NND in the light were shorter than in the dark. In the dark, values were identical to those obtained by simulation of a random distribution of the fish (black bars in Fig. 4B). Thus, SF shoal in the light and light is required for shoaling. This behavioral pattern was markedly different when SF groups were habituated in their tanks for a few days (Fig. Supp. 2): there, SF decreased shoaling in the light and reached IID values similar to CF or random distribution of fish, and their mean IID at night was greater than for a random distribution of fishes, suggesting that they voluntarily moved away from each other.

For CF on the other hand, IID and NND were alike those obtained with a random distribution of fish, showing that they do not shoal. Finally and regarding the *mao* genotypes, SF with or without P106L, and CF with or without P106L were indistinguishable, both in the light and the dark, both without or with habituation (Fig. 4B and Fig. Supp. 2). Therefore, the *mao* P106L mutation has no influence on the evolution of shoaling in *A. mexicanus*.

Next, we studied locomotion, a “simple” trait. At 6dpf, CF larvae were more active than SF larvae, but again there was no influence of the P106L mutation on the distance swam (Fig. 5A). Noteworthy, the locomotor activity of the 2 morphotypes followed day-night variations, but it seemed that only SF anticipated them (at dawn).

For older fish aged 2 or 5 months, CF were still much more active than SF (Fig. 5B). The distance travelled by mutant SF was similar to that of their wildtype siblings; but in CF at these older ages, an effect of the P106L *mao* mutation progressively emerged, with mutant CF being more active than CF carrying the wildtype *mao* allele. Thus, the P106L mutation specifically enhanced locomotion in adult CF, but not in SF.

The locomotion recordings above were performed after 1h of habituation, as commonly done in zebrafish assays. Yet, we noticed that CF placed in a novel tank for the locomotion assay displayed strong thigmotaxis and frenetic swimming, head against walls of the tank (Fig. 5C), i.e., behaviors described as anxiety-related behaviors in zebrafish (Blaser and Gerlai, 2006; Maximino et al., 2010; Schnörr et al., 2012). This occurred immediately after the transfer into the test tank and continued after 1hour (Fig. 5C), suggesting that CF were still stressed by the novel environment. Moreover, SF also still displayed anxiety behaviors such as freezing 1hour after transfer in a novel tank (not shown). We concluded that a 1-hour habituation period was insufficient to quantify accurately locomotion, without being parasitized by stress behaviors. Also, as SF are social animals as shown by schooling, being alone in a tank certainly could be stressful.

To circumvent these biases of the classical locomotion assay, we next studied locomotion in groups and analyzed the time-course of activity during 5 days (Fig. 5D). Strikingly, CF were more active than SF during the first two days, but progressively decreased locomotion and reached SF levels after 60 hours. Moreover, after 3 days of habituation there was no more influence of the morphotype nor of the P106L mutation on the activity level. These data suggested that the P106L *mao* mutation increased locomotion only if the fish was alone and in a novel tank.

Finally, since serotonin plays a role in the control of food intake (Pérez-Maceira et al., 2016; Voigt and Fink, 2015) we quantified food intake in CF, alone or in groups (Fig. 6A). Alone, P106L mutant CF ate twice less food than non-mutant CF. Heterozygotes were intermediate. In groups, fish with the 3 *mao* genotypes ate more than when they were alone, and there was no more influence of the P106L mutation on food intake. Thus, and similar to locomotion, the P106L mutation affected food intake only when the fish were alone.

A mutation in the *mc4r* gene, which causes hyperphagy in CF, has been described (Aspiras et al., 2015). This mutation is not fixed in the Pachón CF population, so we genotyped the individuals used in our food intake assays. All the Pachón fish used (with or without P106L) carried the same *mc4r* allele, the mutant one (Fig. 6B). Thus, the differences in food intake reported above cannot be due to melanocortin signaling, and are specific to the *mao* P106L mutation.

Taken together, these data suggested that *mao* P106L was not responsible for the evolution of aggressive or social behaviors it was previously suspected to control. Importantly, for other traits such as locomotion and feeding, the effects of P106L strongly depended on the context (habituation time, group size), suggesting that these behavioral phenotypes were rather read-outs of another parameter, such as stress or anxiety.

### Effects of the *mao* P106L mutation on stress behaviors, cortisol and catecholamine levels

We measured adrenalin (Adre) and noradrenalin (NA) levels in bodies and heads of 6dpf larvae. Mutants CF had elevated levels of NA in heads (Fig. 7A) and bodies (not shown), and elevated levels of Adre in bodies (Fig. 7A). Importantly, the Adre and NA levels were identical in SF and non-mutant CF, demonstrating that the P106L mutation alone is fully responsible for the increased levels of the two “stress catecholamines” in CF carrying mutant alleles.

**Figure 7.**
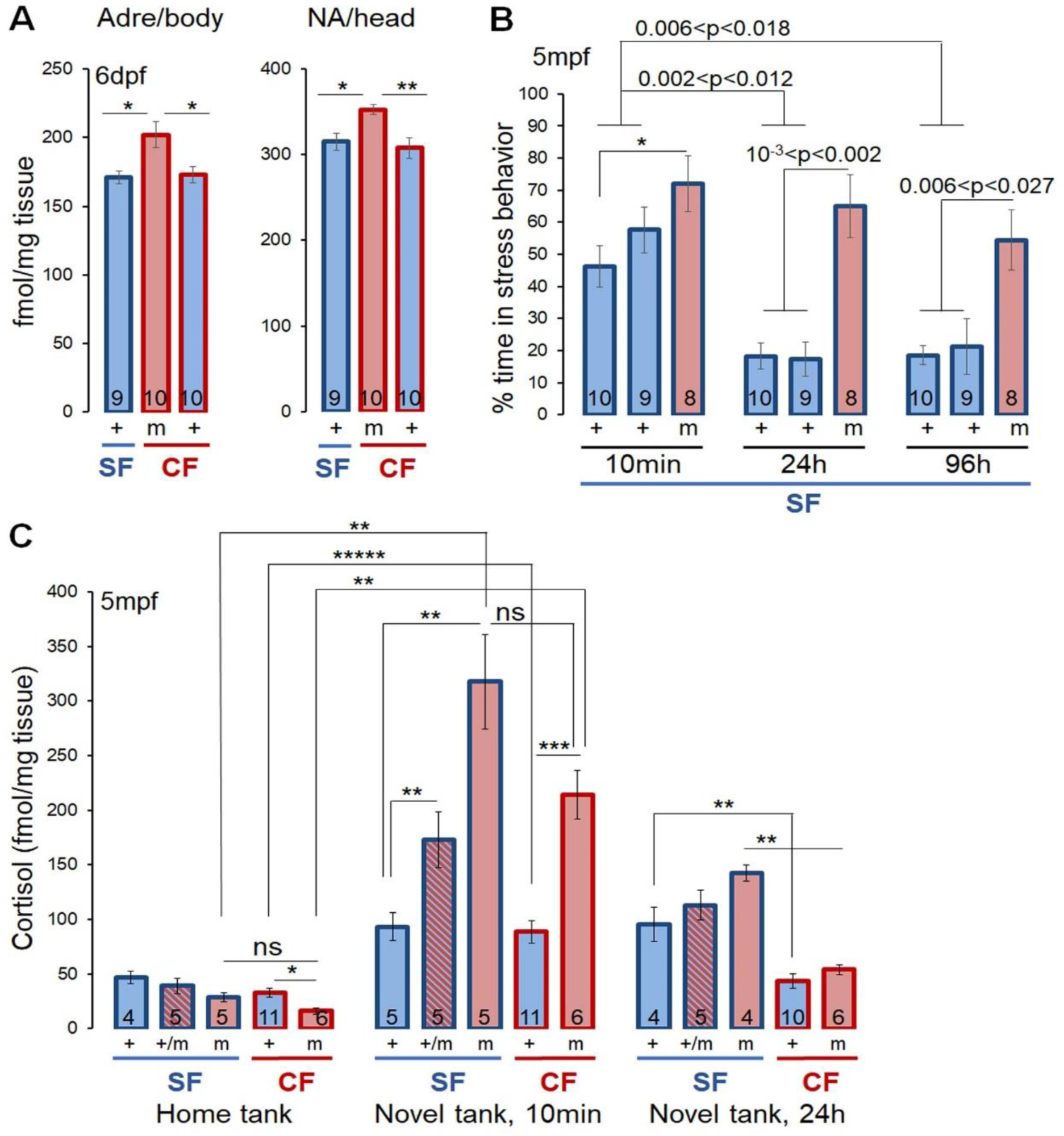
Catecholamine, cortisol and stress behavior measurements. (A) Adrenaline and noradrenaline measurements in the body and the head respectively, of SF, P106L mutant CF (m) and non-mutant CF (+) 6dpf larvae. (B) Percentage of time spent doing stress behavior (freezing, thigmotaxis, erratic movements and attempts to dive) in 5 month-old SF. Fish lines are indicated. Non-mutant SF (+) tested are either wildtype (first bar), or mutants siblings (second bar). Recordings were performed 10min, 24h and 96h after the fish was transferred alone in a novel tank. (C) Cortisol measurements in 5 month-old fish. Measurements were performed on fish held in groups of 6 individuals in their fish facility home tank, in a novel tank for 10 min, and in a novel tank for 24h.

Together, data on locomotion alone in a novel tank, food intake, and catecholamine levels suggested that *mao* P106L could increase the anxiety of the fish. To test this hypothesis, we quantified stereotyped stress behaviors 10min, 24h and 96h after fish were placed alone in a novel tank. For SF, we quantified 4 behaviors: freezing, thigmotaxis, erratic movements, and attempts to dive (see Methods).

A first remarkable observation was that SF had individual preferences for the expression of stress behaviors: some fish almost exclusively displayed freezing, while some others only showed thigmotaxis, and this preference could vary along time (Fig. Supp. 4A-B). In consequence, it was impossible to compare anxiety by quantifying specific stress behaviors independently. Instead, we considered the total time spent showing at least one of these 4 types of stress behaviors (Fig. 7B). At 10min after the beginning of the test, mutant SF and their wildtype siblings spent the same time exhibiting stress behaviors. After 24h and 96h of habituation, mutant SF spent much more time exhibiting a stress behavior than their siblings and wildtype SF. We concluded that the P106L *mao* mutation increased anxiety in SF.

On the other hand, measures of the time performing thigmotaxis in CF (i.e., the only stress behavior expressed by the cave morphs among the 4 defined above) did not show any difference between mutant and non-mutant fish (Fig. Supp. 3 and Video Supp. 1-3).

To bring final support to the involvement of the P106L *mao* mutation in the evolution of anxiety behaviors in *A. mexicanus*, we measured cortisol levels, a reliable indicator of stress, in the 4 fish lines. We measured cortisol 10min and 24h after fish had been placed in a novel tank, as well as in home tanks at the fish facility (Fig. 7C). In their home tanks, P106L mutant CF had less cortisol than non-mutant CF. The same tendency was observed for SF (p=0,063). Thus, the P106L mutation seemed to reduce the basal cortisol levels in a familiar tank. After 10min in a novel tank, cortisol levels increased for all the lines, as expected. For both morphs, the *mao* mutants showed cortisol increases that were about twice higher than their non-mutant counterparts (Fig. 7C). SF heterozygotes were intermediate. This demonstrated that the P106L mutation potentiates the stress reaction in response to a change of environment. Finally, after 24h in a novel tank, cortisol levels were back to initial levels for non-mutant CF, but not for mutant CF and for the 3 types of SF. Moreover, mutant SF still had slightly higher cortisol level than non-mutant SF (p=0.057). Altogether, this suggested that the mutation still influenced cortisol/stress, even after 24h in a novel tank: P106L mutants needed a longer time to habituate. Finally and regarding the SF/CF comparison, we found that in CF cortisol levels went back to baseline quicker than SF. Thus, we have uncovered an interaction between the P106L *mao* mutation, which confers stressability in *A. mexicanus*, and a morph-dependent variable, which increases the time of recovery to baseline in SF.

Altogether, these results showed that the P106L *mao* mutation increases anxiety in a novel environment in *A. mexicanus*.

### Effects of P106L mutation on other candidate phenotypes

In mice, serotonin and MAO-A have an effect on cardiac rhythm, development and function (Abzalov et al., 2015; Mialet-Perez et al., 2018; Nebigil and Maroteaux, 2001; Nebigil et al., 2000; Stoyek et al., 2017). Yet, CF have a low metabolic rate (Hüppop, 1986; Moran et al., 2014; Salin et al., 2010), so we checked if *mao* P106L may modify cardiac rhythm in *A. mexicanus*. In both morphs, mutant fish presented the same cardiac rhythm as their non-mutant counterparts (Fig. 8A), suggesting no influence of the P106L mutation on this physiological parameter.

**Figure 8.**
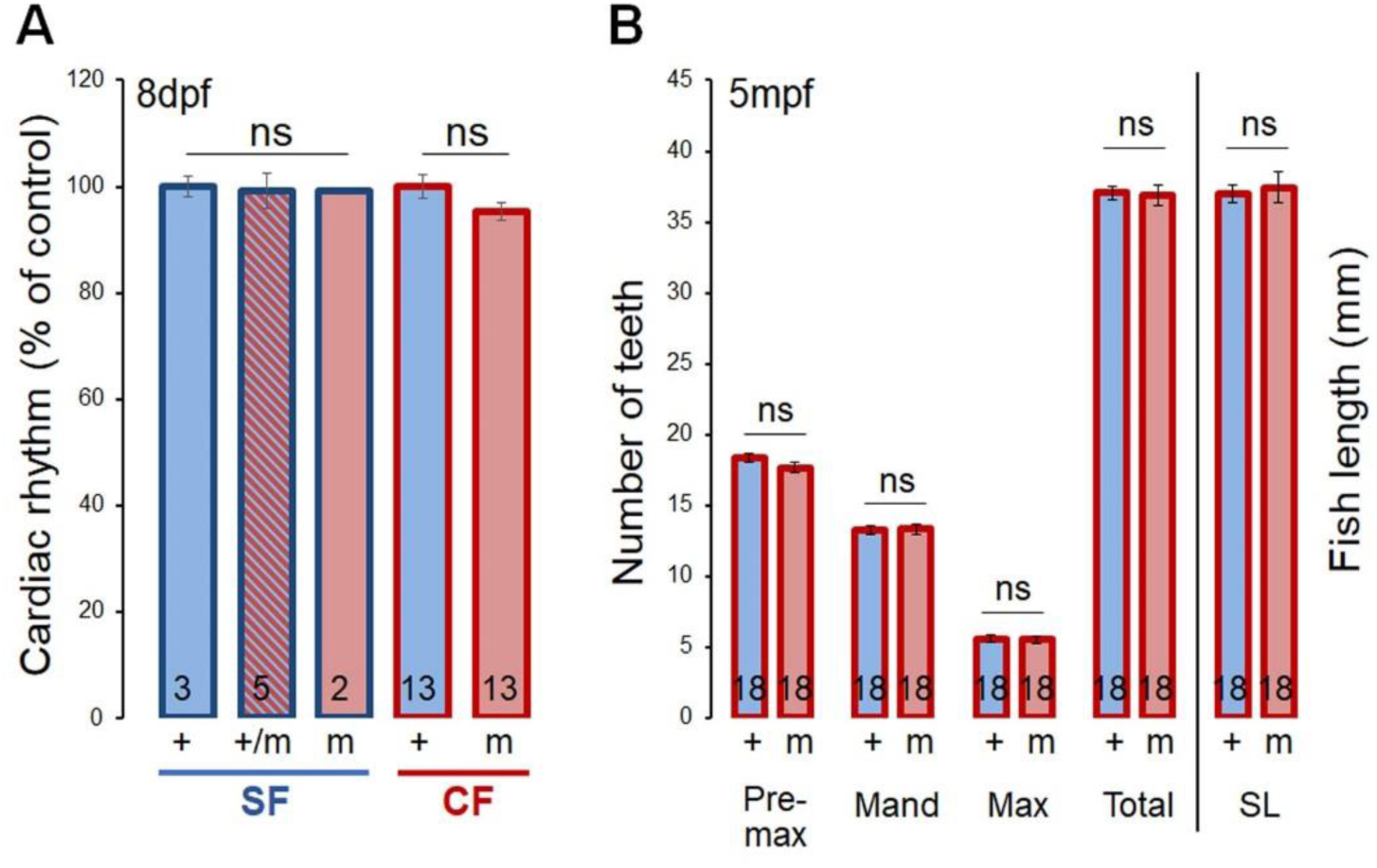
Cardiac rhythm and number of teeth. (A) Cardiac rhythm of 8dpf larvae, in the indicated fish lines. (B) Number of teeth on pre-maxilla, mandible and maxilla, total number of teeth, and fish length in P106L mutant (m) and non-mutant (+) CF.

Serotonin also plays a role on the craniofacial skeleton and teeth development (Moiseiwitsch, 2000; Moiseiwitsch et al., 1998; Reisoli et al., 2010). Yet, CF possess more teeth than SF (Atukorala et al., 2013), so we tested if *mao* P106L could increase the number of teeth. Mutant CF had the same number of teeth as non-mutants, also ruling out any influence of the P106L mutation on this anatomical trait, nor on fish size (Fig. 8B).

## Discussion

### *mao* alleles in the wild

Through sampling in wild populations of *A. mexicanus*, we built a phylogeographic map of *mao* alleles. The P106L mutation was present exclusively in El Abra caves, with the mutant allele found in large proportions, or even fixed in two caves. Thus, either the P106L mutation has been selected, or the observed proportions have been reached by genetic drift, possibly helped by bottleneck effects at the time of colonization or later.

Concerning the origin of the P106L mutation, we can imagine two hypotheses. (i) the mutation appeared in a cave of the El Abra group, and then spread to other caves through karstic subterranean water networks, or (ii) the mutation appeared in the ancestral surface population, and individual(s) carrying the mutation entered one or several caves of El Abra. According to this last hypothesis, the P106L mutation may still be present in the surface population, in low proportion, as it is the case for the *mc4R* mutation (Aspiras et al., 2015). As we did not find the mutated allele in the river populations, we could not discriminate between the two hypotheses from our dataset.

The fact that the mutation is not fixed in most El Abra caves suggests that it is recent. In a further attempt to reconstruct the evolutionary history of the mutation, we screened all polymorphisms in a region of ∼4kb around the P106L mutation, for SF and for CF originating from nine different caves, in 110 wild-sampled individuals (Fig. Supp. 6). The goal was to find haplotypes corresponding to the mutant and the wildtype *mao* allele, respectively. The total number of polymorphisms encountered in the 4kb *mao* fragment was consistent with what is known about the ecology of different populations (Fig. Supp. 6A): polymorphism was high in the surface population, which is very large, as well as in the Chica cave population, where the introgression of surface alleles is frequent, as witnessed by the presence of SF and hybrids in the pools (Elliott, 2018; Mitchell et al., 1977; Torres-Paz et al., 2018). Of note, this also explains why the *mao* P106L allele is found at lower frequency in Chica compared to other El Abra caves. Conversely, polymorphism was lowest in the Curva cave – where the P106L mutation is also fixed. The Curva cave is tiny, and even if no estimation of population size is available, we can suppose it is very small. Our data regarding shared polymorphisms between cave populations are also consistent, with many polymorphisms shared inside groups of caves, and few between groups (Fig. Supp. 6B). Unfortunately, even if clear haplotypes emerged (Fig. Supp. 6C), we could not propose a simple scenario for the diffusion and evolutionary history of the P106L mutation in caves, since too many recombination events happened. Nevertheless, haplotype distributions strongly suggest that the P106L mutation has travelled and colonized El Abra caves without being counter-selected (conclusions on positive selection cannot be drawn, see below).

Among the SF individuals, we have incidentally found a non-synonymous mutation just next to the P106L position: M107I. This mutation has a MutPred2 score of 0.247, which is the probability that the amino acid change is pathogenic (Pejaver et al., 2017). By comparison, the P106L mutation has a MutPred2 score of 0.456, close to the pathogenicity threshold (0.5). Therefore, the M107I mutation found in surface is probably not deleterious for the enzyme function.

### Cavefish behavioral syndrome

Many studies have demonstrated the involvement of serotonin in aggression in vertebrates (Edwards and Kravitz, 1997; Nelson and Trainor, 2007; Olivier, 2004; Popova, 2006). In most animal models, an increase in 5-HT neurotransmission causes a decrease in aggression (Carrillo et al., 2009; Summers and Winberg, 2006; Winberg and Nilsson, 1993). Indeed, pharmacological treatments with selective serotonin reuptake inhibitors (SSRIs) as fluoxetine/Prozac or serotonin receptors agonists generally decrease aggression, whereas serotonin receptors antagonists or the inhibitor of serotonin synthesis pCPA generally increase aggression (Buchanan et al., 1994; Deckel and Fuqua, 1998; Filby et al., 2010; Lopez-Mendoza et al., 1998; Lynn et al., 2007; Perreault et al., 2003; Sperry et al., 2003). Moreover, long-term supplementation in tryptophan (5-HT precursor, an essential amino acid) in food regime also causes an activation of 5-HT system, and reduces aggression in trout (Winberg et al., 2001). However, some studies also found no changes of aggression after manipulations of 5-HT neurotransmission (Filby et al., 2010; Winberg and Thörnqvist, 2016), suggesting a complex role for serotonin in the expression of agonistic behavior in fish.

In the species *Astyanax mexicanus*, cavefish are less aggressive than surface fish (Elipot et al., 2013; Espinasa et al., 2005). Therefore, the P106L *mao* mutation was an ideal candidate to serve as the genetic basis of reduced aggression. However, here we demonstrate that even though *mao* P106L causes an increase in 5-HT levels, it does not affect aggressive behavior, regardless of vision being present or not. It is even more surprising because Elipot et al. (2013) had shown that SF under acute fluoxetine treatment were less aggressive. A possible explanation is that an acute increase in 5-HT produced pharmacologically may be very different from the chronic inhibition of MAO activity produced by the mutation, which could induce plastic changes in neuronal networks and homeostatic compensations to elevated 5-HT levels. Elipot et al. (2013) had also performed developmental manipulations of the 5-HT system in CF using cyclopamine (a Sonic Hedgehog signaling pathway inhibitor), which resulted in a decrease in size of the PVA and raphe neuronal clusters and an increase in aggression. Here, we did not observe any change in size of 5-HT clusters caused by the P106L mutation; only changes in 5-HT levels, which may not suffice to change aggression. Finally, as dopamine (DA) has a stimulatory role on aggression in mammals (Nelson and Trainor, 2007; Rodriguiz et al., 2004) and maybe also in fish (Filby et al., 2010; Winberg and Nilsson, 1993), we can hypothesize that the increase in both 5-HT and DA levels caused by the P106L *mao* mutation could compensate each other, resulting in no change in aggression.

Schooling or shoaling (as studied here) has advantages as it reduces the threat of predation and facilitates visual food research (Miller and Gerlai, 2011). Schooling behavior is supported by vision and lateral line (Partridge and Pitcher, 1980). Here, we have shown that light is necessary for SF to shoal, which is consistent with vision being required for this behavior (Kowalko et al., 2013). We have also shown that SF shoal less when they are in their tank since few days, consistent with other studies describing a reduction of shoaling due to the habituation to the environment (Al-Imari and Gerlai, 2008; Delaney M. et al., 2002). Indeed, fear is one of the driving force of shoaling, and if habituated fish are disturbed, they start schooling again (personal observation).

To form a school is important in predator detection and escape. But it requires vision, and becomes dispensable in environments such as *Astyanax* caves where predators are absent. Moreover, in the dark, anti-predator advantages of forming a school are lost, and schooling may be disadvantageous since from the predator point of view, all preys are in the same area. Most of the social fish species do not school at night (Pavlov and Kasumyan, 2000). We found that shoals of *A. mexicanus* SF disperse at night, too.

Little is known about neural bases of schooling, even if some studies suggested the implication of the DA system. Indeed, treatments with DA agonists or antagonists affect schooling (Echevarria et al., 2008; Scerbina et al., 2012). Even though the P106L mutation changes brain DA levels, we could not show any effect on schooling, again regardless of the fish having eyes or not. Finally, Kowalko et al. (2013) showed reduced schooling in SF after deprenyl treatment (MAO inhibitor). As discussed above, a possible explanation is that deprenyl treatment leads to an acute inhibition of MAO activity, whereas the mutation corresponds to a chronic inhibition.

Previous studies on *A. mexicanus* have shown that CF are more active than SF at adult (Carlson and Gross, 2018; Yoshizawa et al., 2015) and larval (Duboué et al., 2011) stages. In larvae this difference is due to both reduction of sleep and increased waking velocity in CF (Duboué et al., 2011). In adults, it is mostly due to the reduction of sleep, and hyperactivity has little contribution to the phenotype (Yoshizawa et al., 2015). Our results are consistent with that: we showed a higher locomotor activity in larval CF, but no difference in adults from 72h of habituation. Of note, we did not measure sleep, so we cannot know the parts of sleep loss and waking velocity in these results. In another study on adults, Carlson et al. (2018) showed higher activity in CF than in SF. A major difference with our study is that they recorded single fish whereas we recorded groups of fish. SF are social animals, and a SF put alone in a tank displays more freezing than in a group (personal observation). It is therefore not surprising to observe reduced locomotion in SF compared to CF in solo condition.

Importantly, it appears that 1h is not enough for habituation to a novel environment in our fish model, as both SF and CF still display stress behaviors. The difference of locomotion between CF and SF (alone or in groups) measured 1h after transfer in the test tank could be explained by the fact that CF spent almost 100% of their time doing thigmotaxis (hence they move), whereas SF also spent some time freezing (hence no movement). In these conditions, it is therefore doubtful that the activity recorded is “true” locomotion. It rather corresponds to a biased read-out of stress behaviors. In line with this remark, the only effect of the *mao* mutation we observed on “locomotion” was for 5months old CF put alone and in a novel tank since 1h (Fig. 5B, right). Both mutant and non-mutant CF spent almost 100% of their time doing thigmotaxis, so we can conclude that the difference in distance travelled is only due to increased velocity in mutants. The swimming of mutant CF was more frenetic, which is consistent with our measures of cortisol levels. We did not observe this difference in frenzy/velocity when the fish were in groups. This suggests that being in a group has some anxiolytic effect, even in CF, usually considered as non-social animals.

Many studies have reported a decrease of locomotor activity when serotonin levels are increased (Fingerman, 1976; Gabriel et al., 2009; Perreault et al., 2003; Winberg et al., 1993). Conversely, an increase of DA activity usually results in increased locomotion in mammals (Beninger, 1983) and fishes (Boehmler et al., 2007; Bretaud et al., 2004; Godoy et al., 2015; Irons et al., 2013; Jay et al., 2015; Lambert et al., 2012; Tran et al., 2015). However, although the P106L mutation changes 5-HT and DA levels, it did not affect locomotor activity. We may hypothesize that the changes in 5-HT and DA levels compensate.

Several pieces of evidence converge towards the idea that P106L *mao* mutation increases fish anxiety: higher catecholamine levels, lower food intake when fish are alone in their tank, and higher locomotion in a novel tank. The literature also supports a link between serotonin and anxiety (Chen et al., 2004; Herculano and Maximino, 2014; Lillesaar, 2011).

To validate this hypothesis, we quantified some stress behaviors using a novel tank test, known to induce anxiety in fishes (Bencan et al., 2009; Blaser et al., 2010; Cachat et al., 2010; Egan et al., 2009; Kysil et al., 2017; Levin et al., 2007) and other species (Moriarty, 1995; Simon et al., 1994; Treit and Fundytus, 1988). Fish have many displays of stress behaviors which are used to measure anxiety: freezing, thigmotaxis (also called wall-following, thrashing or escape), erratic movement, leaping, bottom dwelling (or diving) (Bencan et al., 2009; Blaser and Gerlai, 2006; Blaser et al., 2010; Cachat et al., 2010; Egan et al., 2009; Gerlai et al., 2006; Levin et al., 2007; Maximino et al., 2010; Schnörr et al., 2012; Speedie and Gerlai, 2008). *Astyanax* SF display all of these behaviors in a novel tank (Chin et al., 2018; personal observations), whereas CF only do thigmotaxis (Abdel-Latif et al., 1990; Patton et al., 2010; Riedel, 1998; Sharma et al., 2009; Teyke, 1989). Such difference in stress-elicited behaviors could be due to the absence of predators in natural cave environment. In rivers, stress behaviors like freezing, erratic movements and diving are necessary to escape from predators and hide in plants or rocks in the river bottom. Whereas in caves, these behaviors are probably not selected, and maybe even counter-selected, since freezing reduces the time spent for foraging, erratic movement is energy consuming, and food can fall and stay at the surface of water, so bottom dwelling is also probably non-adaptive.

Consistent with our hypothesis, we showed that P106L mutant SF spent more time expressing anxiety behaviors than non-mutant SF. Also, we expected more thigmotaxis in mutant CF than in non-mutants, but it turned out not to be the case. A possible explanation could be that in CF, the robust thigmotaxis elicited by a novel environment may not entirely be an anxiety behavior and an attempt to escape, but also a form of exploratory behavior.

In zebrafish and rodents, thigmotaxis is considered an anxiety behavior, since anxiolytic treatment like diazepam decrease it, and anxiogenic treatment like caffeine promotes it (Schnörr et al., 2012; Treit and Fundytus, 1988). Such experiments were not conducted in *Astyanax* SF, but the behaviors of SF in a novel tank are very similar to those of zebrafish. Cavefish may be different. Interestingly, in CF the duration of thigmotaxis is longer in a complex environment than in a simple environment (Teyke 1989). The duration of thigmotaxis is also shorter when the fish is placed in a familiar environment, but not when the consolidation of the memory of that environment has been impaired. Moreover, CF can detect spatial changes in their environment (Burt De Perera, 2004), and fish can built a cognitive spatial map (Burt De Perera, 2004; Rodriguez et al., 1994). Finally, it seems like CF swim slightly more parallel to the wall than SF when they do thigmotaxis (Sharma et al., 2009), consistent with the fact that thigmotaxis could correspond to a different expression in CF (exploratory) and SF (escape). On the other hand, spatial learning in fishes involve the telencephalon, particularly the pallium (Broglio et al., 2010; Rodríguez et al., 2002; Saito and Watanabe, 2006), and CF still display thigmotaxis and habituation after telencephalic ablation (Riedel, 1998). Moreover, since thigmotaxis is an anxiety behavior in SF and many other species, including humans (Kallai et al., 2005; Kallai et al., 2007), the most parsimonious hypothesis is that it is also an anxiety behavior in CF. A simple way to confirm (or not) this idea in the future will be to treat CF with anxiolytic or anxiogenic drugs. Finally, it may be that the two hypotheses (anxiety or exploratory) are not exclusive, and thigmotaxis elicited by stress could permit to explore the environment. In any case, there must be a stress component in thigmotaxis behavior, as CF placed in a novel environment do show marked increase in cortisol levels.

In fish (as in mammals), an increase of plasmatic cortisol levels occurs after or during a stressor, like predator visuals (Barcellos et al., 2007), crowded environment (Ramsay et al., 2006), social stress (Øverli et al., 1999; Tea et al., 2019), chasing (Gesto et al., 2008), confinement (Backström et al., 2011; Gallo and Jeffery, 2012; Kiilerich et al., 2018; Schjolden et al., 2006; Vijayan et al., 1997), or novel tank (Kysil et al., 2017). Cortisol levels are commonly used as stress response indicator. During or after a stressor, the hypothalamus releases corticotropin-releasing factor, which induces the release of adenocorticotropic hormone (ACTH) in the blood stream. ACTH in turn stimulates the release of cortisol by interrenal cells of the head kidney. This is called the hypothalamic-pituitary-interrenal axis (HPI axis), homologue to the hypothalamic-pituitary-adrenal (HPA) axis of mammals (Mommsen et al., 1999; Wendelaar, 1997). Cortisol has multiple effects on organs and tissues. For instance, it induces a reallocation of energy to cope with stress, with an increase of gluconeogenesis, lipolysis, proteolysis, and an inhibition of growth, reproduction, and immune responses (Aluru and Vijayan, 2009; Faught and Vijayan, 2016; Harris and Bird, 2000; Mommsen et al., 1999).

Previous studies concluded to a reduction of anxiety in CF as compared to SF (Chin et al., 2018), but here we showed that the difference between the two morphs is more subtle than “more or less stressed”. In normal conditions (home tank in fish facility), P106L mutants have lower levels of cortisol than non-mutants. In a novel tank, P106L mutants have higher cortisol levels. In the wild, since most fish in the Pachón cave are mutants, we can assume that they are less stressed than SF in normal conditions, and more stressed in a stressful condition. An explanation to these results may lie in the link between monoamines and cortisol release during stress response. Indeed, positive or negative modulations of cortisol release by catecholamines and serotonin are known (Rotllant et al., 2006; Medeiros et al., 2010; Winberg et al., 1997; Höglund et al., 2002; Saphier et al., 1995), and this would regulate the intensity of the stress response (Gesto et al., 2015). Here, 10min after stress we detected a 2-fold variation of brain NA and 5-HT levels for mutant fish but not for wildtype ones, and in both morphs (Fig. Supp. 5 and its legend), providing further support to the involvement of monoamines in the regulation of stress response.

We suggest that the change in basal monoamine levels due to the P106L mutation could alter the balance of neuromodulation (5-HT and DA) on the HPI axis. It is also possible that chronic inhibition of MAO by the mutation could induce neuronal plasticity and changes in the monoamine network, and their effect on the HPI axis. Indeed, an acute inhibition of MAO by deprenyl treatment has no effect on cortisol levels in human (Koulu et al., 1989), while MAO-A knockout/inactivation induces a decrease in corticosterone levels (Popova et al., 2006). Why a chronic inhibition of MAO induces changes in the regulation of HPI axis remains unclear and needs further investigations.

Finally, we have shown that cortisol levels of CF return to basal levels quicker than those of SF. Therefore, there is a morph-dependent parameter which accelerates the decrease of cortisol level in CF. Strickler and Soares (2011) have shown that cannabinoid receptor *CB1* expression is upregulated in CF, and the endocannabinoid system is known to modulate the HPA stress axis in mammals, specifically to enhance recovery to baseline following stress (Hillard et al., 2017; Micale and Drago, 2018). The upregulation of *CB1* in CF could be a good candidate to explain faster recovery of basal cortisol levels in CF.

### *mao* P106L: selected or not?

The high proportion of mutant alleles in El Abra caves makes us wonder if the mutation could be advantageous for life in caves, and may be selected. Conversely, we can suppose that it would be counter-selected in rivers.

Epigean rivers appear like stressful environments, with both biotic (predators, parasites) and abiotic stressors (changes in temperature, salinity, turbidity, water levels). Conversely, caves are more stable/buffered environments, with no predators, probably less parasitism (intermediate hosts are not always present in caves), and little changes of temperature, salinity, turbidity and water levels (Mitchell 1977). Because of the deleterious effects of cortisol on growth, immune response, reproduction and metabolic rate, chronic high levels of cortisol are susceptible to impair all these physiological processes (Madison et al., 2015; Pickering and Pottinger, 1989).

For CF, having very low levels of cortisol in standard condition could be adaptive since the growth, reproduction and immune system would be spared. Moreover, this could facilitate energy storage and participate to the lower metabolic rate measured in CF (Hüppop, 1986; Moran et al., 2014; Salin et al., 2010). Since caves are a “quiet” environment, at least during the dry season, we can suppose that the wide increase of cortisol levels observed with a stressor in our lab study would not occur frequently in the wild – and in any case, they would rapidly return to basal levels, as we have shown.

For SF on the other hand, lower basal levels of cortisol could decrease alertness, and impair energy mobilization and predator escape. Moreover, higher cortisol levels induced by a moderate stress, and so, overreaction to that stress, could be deleterious in the long term. This is consistent with Brown et al. (2005) who showed that *Brachyraphis episcopi* from high-predation environment have a lower stress response to a stressor than individuals from low-predation environment. This could prevent energy expenditure for small stressors, and save energy for important stressors like predators.

Thus, the mutation may actually be counter-selected in surface and positively-selected in caves. A less “adaptationist” hypothesis would be that the mutation is essentially neutral in caves, and evolves under a genetic drift regime. Further studies will need to address this question. Of note, since monoaminergic systems are involved in a variety of developmental, physiological and behavioral mechanisms, it is possible as well that the P106L *mao* mutation has other phenotypic effects, beyond those we have investigated here; and if so, we cannot exclude that these unknown phenotypic effect(s) may contribute to all or part of the selective process, if any.

## Conclusion

The P106L mutation in cavefish *mao* has a significant effect on stress response and anxiety-related behaviors. It is very likely that this effect is responsible for changes in food intake (Conde-Sieira et al., 2018; Wendelaar, 1997) or the “frenzy” observed in a novel tank. The relatively low impact of the mutation on anatomy and the global behavioral syndrome of cavefish is surprising, considering the implication of monoaminergic systems in many physiological, behavioral and developmental processes. The existence of compensatory mechanisms is likely. Indeed, 5-HT and DA seem to have opposite effects on aggressiveness and locomotion, and the P106L mutation increase both 5-HT and DA levels. We can also predict that chronic inhibition of MAO should induce modifications of expression of monoamine receptors, transporters, and other signaling components that rescue a “normal” neuromodulation despite high levels of neurotransmitters. Such compensation phenomena have been described in the literature (Evrard et al., 2002).

## Acknowledgements

Work supported by an Equipe FRM grant (Fondation pour la Recherche Médicale DEQ20150331745) and CNRS to SR. CP received PhD fellowships from the French Ministry of Research and from FRM. We thank Luis Espinasa, Ramses Miranda, Angeles Verde, Ulises Ribero, María de Lourdes Vazquez, Carlos Pedraza, Laurent Legendre, Didier Casane, and all past and present members of the Rétaux’s lab for their help in data collection in the field. We are grateful to Stéphane Père, Victor Simon and Krystel Saroul for taking care of our *Astyanax* colony; Maryline Blin, Victor Simon and Jorge Torres-Paz for technical advices; and Maxime Policarpo for help with genomic databases.

