## Supplemental data file for "A mutation in monoamine oxidase (MAO) affects the evolution of stress behavior in the blind cavefish *Astyanax mexicanus*"

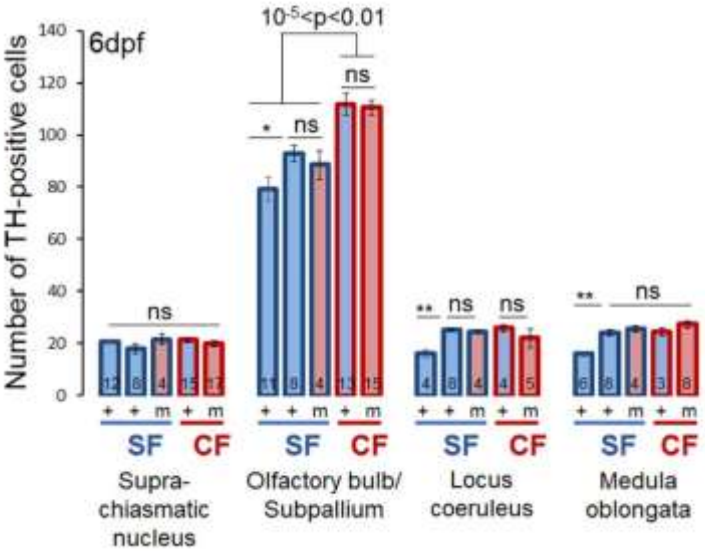


Supp 1

**Figure Supp 1. TH positive cells, additional clusters.**

Number of TH-positive cells in suprachasmatic nucleus, olfactory bulb and subpallium, locus coeruleus and medulla oblongata, for SF and CF, both mutant homozygotes (m) or non-mutant (+). Non-mutant SF (+) tested are either wildtype (first bar), or mutants siblings (second bar).

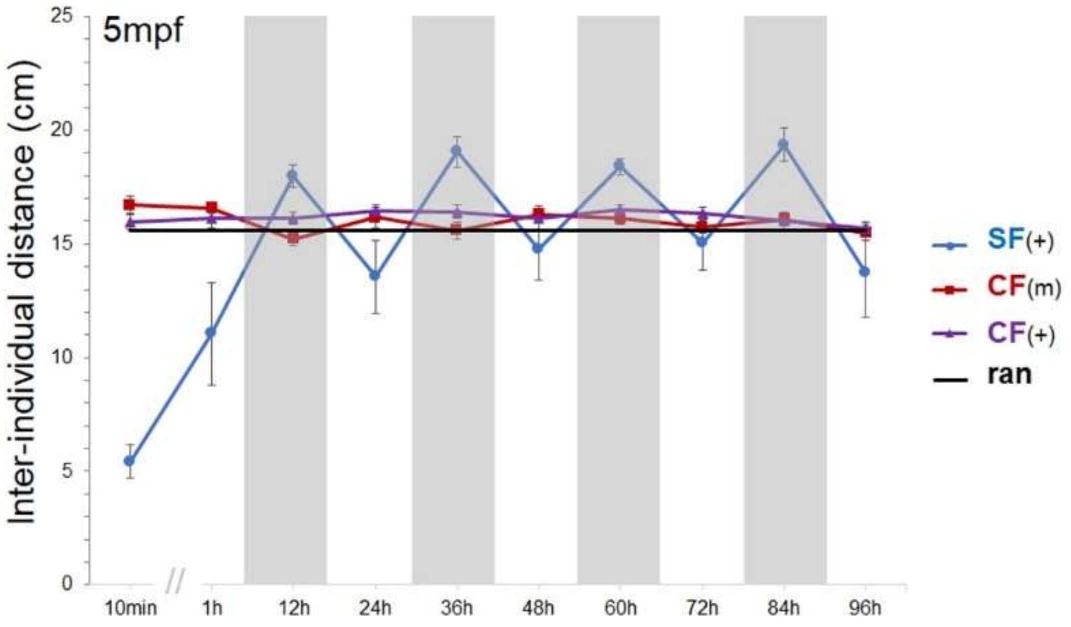


Supp2

**Figure Supp 2. Shoaling in SF and CF, measured over a one week period.**

Mean of inter-individual distance (IID) measured during the day or during the night, over 5 days in groups of 6 fish: n=10 groups for P106L mutant CF (red line), n=12 groups for non-mutant CF (purple line), and n=5 groups for non-mutant SF (blue line). The black line is the mean value of inter-individual distance obtained with 100.000 simulations of random distribution of fish (ran).

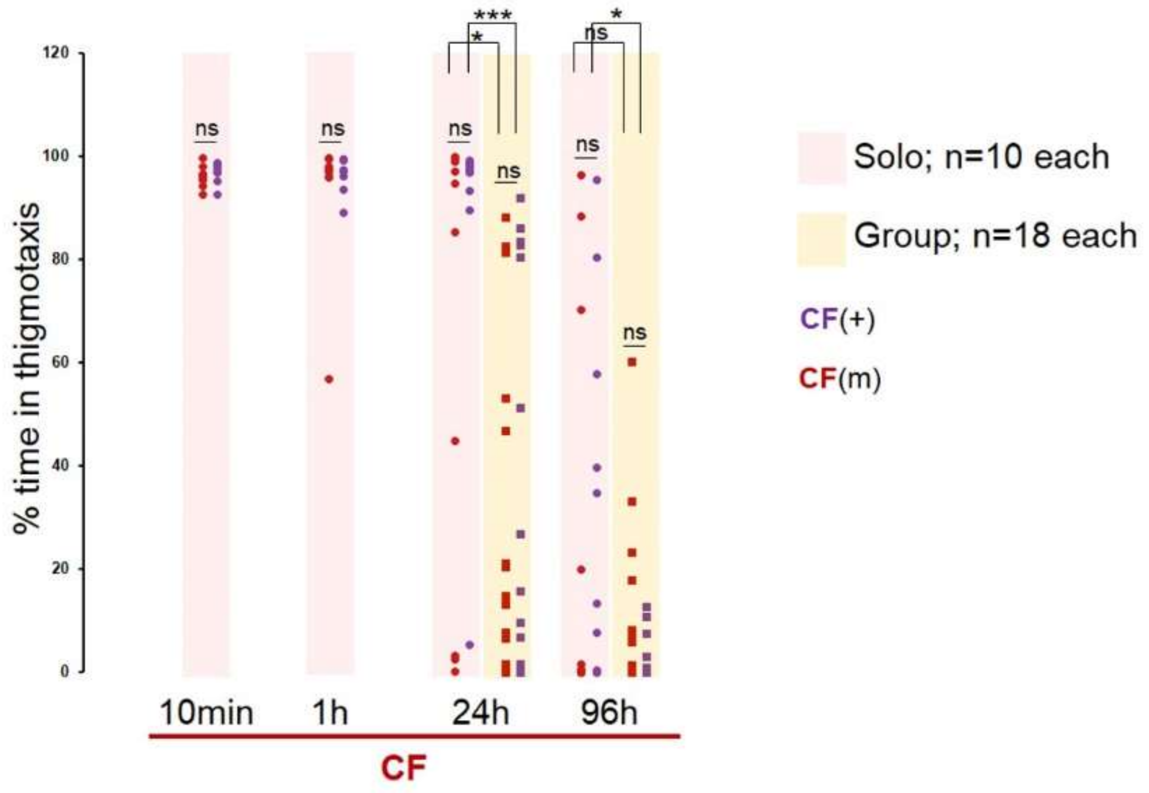


Supp3

**Figure Supp 3. Thigmotaxis in CF.**

Measurements of the percentage of time doing thigmotaxis 10min, 1h, 24h or 96h after the fish was/were put alone (pink shade) or in group (yellow shade) in a novel environment. The measurements were performed on P106L mutant CF (red) and non-mutant CF (purple). Data are presented as scattered plots to represent the inter-individual variability of this phenotype at the two later time-points.

At 10min and 1h after transfer in a new tank, CF spent ~100% of their time doing thigmotaxis, regardless of their *mao* genotype. It was impossible to score groups of fish at these times (i.e., no yellow bars for 10min and 1h) because frenetic swimming led to loss of the track of individual fish (manual scoring).

At 24h and 96h after transfer in a new tank, data could be obtained for the 4 conditions: mutant and non-mutant, solo or groups. As inter-individual variability was high, there was no detectable significant effect of the *mao* mutation on thigmotaxis behavior, either alone or in group. However and interestingly, there was a significant reduction of thigmotaxis, for both the mutant and the non-mutant CF, when the fish were in groups as compared to solo. This may suggest that the group has an anxiolytic effect on CF, although the CF morphotype of *A. mexicanus* is often described as a “non-social” animal because it does not show collective behavior such as schooling or shoaling. Of note, we cannot rule out the possibility that the “anxiolytic effect” observed here was not due to the size of the tank. Indeed, groups were tested in larger tanks (40x23cm; 5l) than single fish (19x10cm; 600ml), hence the possibility of a confinement stress in the later.

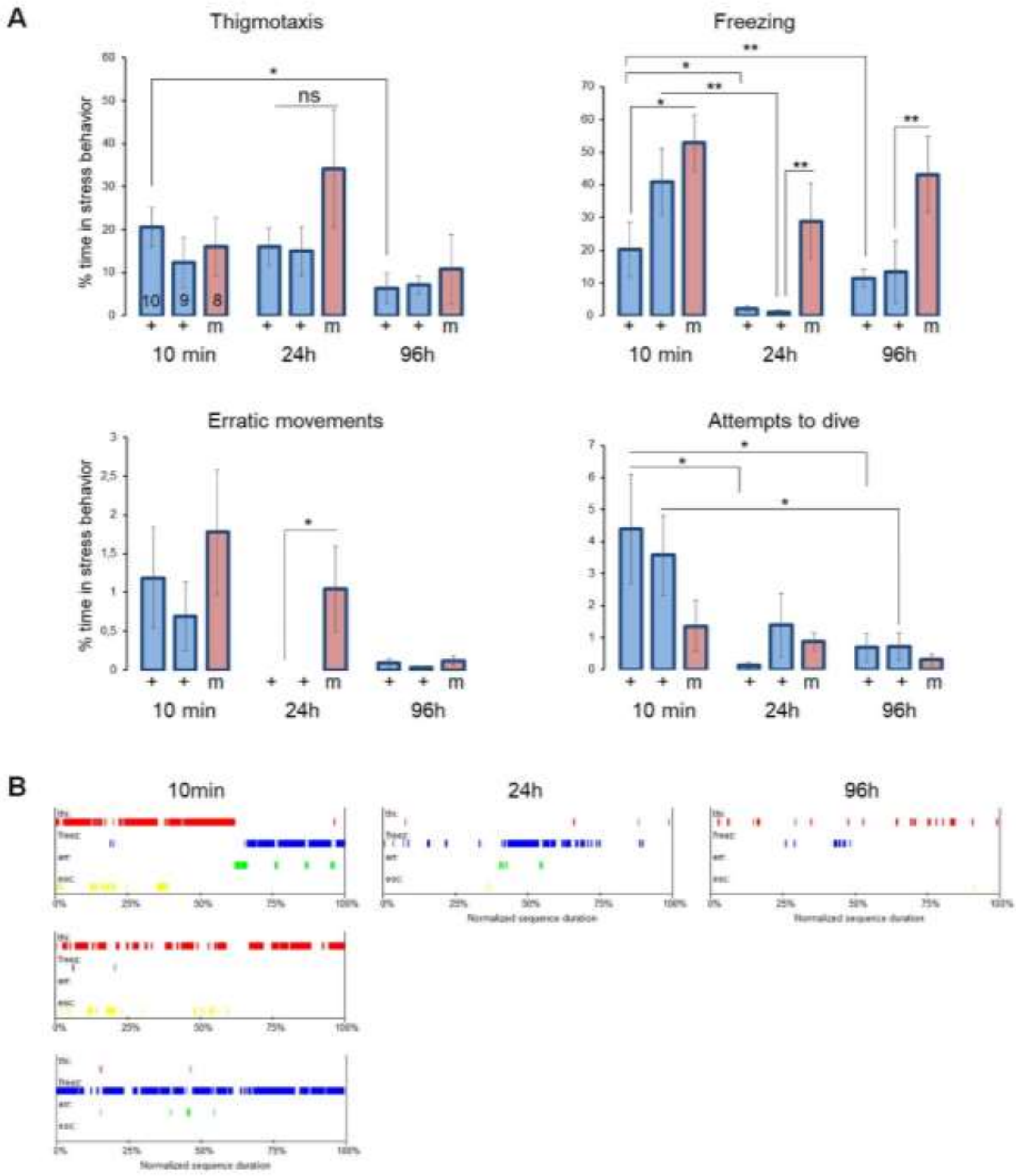


Supp 4

**Figure Supp 4. Stress behaviors in SF.**

(A) Measurements of the percentage of time doing thigmotaxis, freezing, erratic movements and attempts to dive. The measurements were performed on P106L mutant SF (m) and non-mutant SF (+), and 10min, 24h or 96h after the fish was put alone in a novel environment. Non-mutant SF tested were either wt (first bar of each group), or siblings of mutants SF (second bar). (B) Representative ethograms for SF, showing the alternation of periods where the fish display thigmotaxis ('thi' in red), freezing ('freez' in blue), erratic movements ('err' in green) and attempts to dive ('esc' in yellow). The 3 ethograms of the top line belong to the same fish, recorded 10min, 24h and 96h after being put in a novel environment. The 3 ethograms of the first column belong to 3 different fish recorded 10min after being put in a novel environment. Note the individual preferences for a given behavior, which can also vary along time for a single individual. Hence, the large error bars on graphs in A, and the necessity to combine all stress behaviors for analysis (as shown in Fig. 7B).

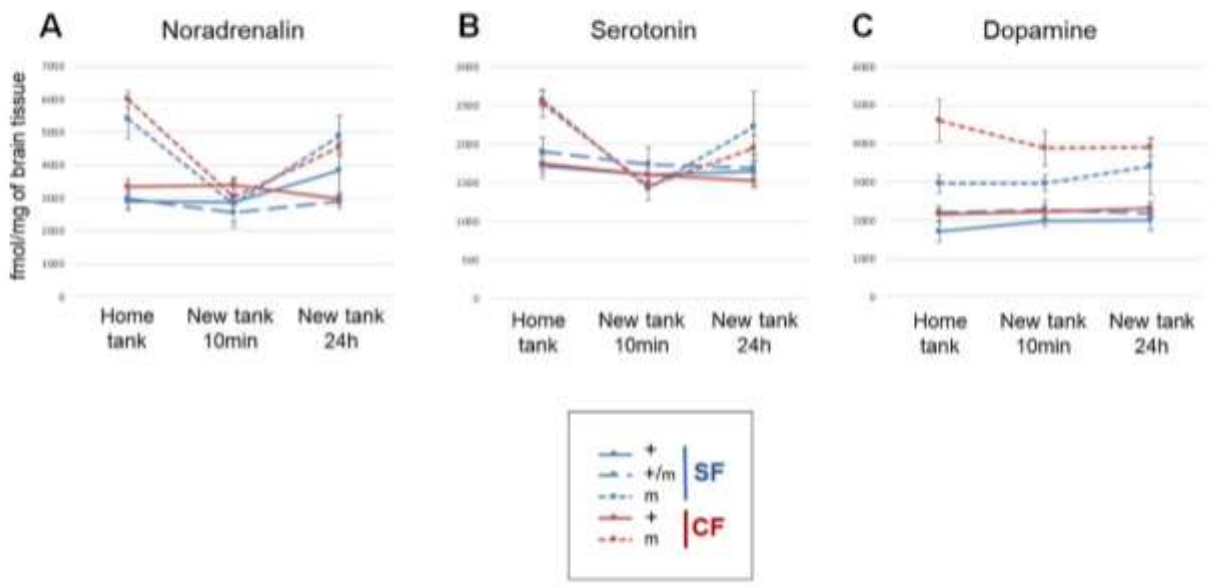


Supp 5

**Figure Supp 5. Levels of serotonin, noradrenaline and dopamine in the brain before, during and after a stressor (novel environment).**

Noradrenaline (A), serotonin (B) and dopamine (C) measurements in the brains SF and CF, both P106L mutant homozygotes (m, dotted lines) and non-mutant (+, continuous lines), and heterozygotes SF (+/m, long dotted lines). Measurements were performed on fish held in groups of 6 in their home tank (HT) at the fish facility, in a novel tank for 10 min (NT10), and in a novel tank for 24h (NT24). Only mutants show strong variations of brain NA and 5-HT after stress.

Statistics for 5-HT: HT mCF versus HT +CF p=0.0011 ; HT mSF versus HT +SF p=0.0159 ; HT mCF versus NT10 mCF p=0.0022 ; HT mSF versus NT10 mSF p=0.0079 ; NT24 mCF versus NT24 +CF p =0.0312.

Statistics for NA: HT mSF versus HT +SF p=0.0159 ; HT mSF versus HT m/+SF p=0.0079 ; HT mCF versus HT +CF p=0.0002 ; HT mCF versus NT10 mCF p=0.0022 ; HT mCF versus NT24 mCF p=0.0022 ; NT10 mCF versus NT24 mCF p=0.0022 ; NT24 mSF versus NT24 m/+SF p=0.0159 ; NT24 mCF versus NT24 +CF p=0.0002.

Statistics for DA: HT mSF versus HT +SF p=0.0317 ; HT mSF versus HT m/+SF p=0.0079 ; HT mCF versus HT +CF p=0.0031 ; NT10 mSF versus NT10 +SF p=0.0159 ; NT10 mCF versus NT10 +CF p=0.0002 ; NT24 mCF versus NT24 +CF p=0.0010.

Comments:

*Noradrenaline*: A stressor induces a rapid increase of noradrenergic activity (Øverli et al., 1999; Øverli et al., 2001; Weber et al., 2012), with activation of the cholinergic sympathetic pathway, which induces catecholamines release by chromaffin cells in the head kidney (Wendelaar, 1997). A paracrine activation of interrenal cells by catecholamines and thus an activation of cortisol release have been shown in fish (Rotllant et al., 2006). Here, 10min after stress we detected a 2-fold variation of brain NA levels for mutant fish but not for wild type ones, both in SF and CF (Fig. Supp. 5A).

*Serotonin*: There is a rapid increase of serotonergic activity during or after a stressor, associated with an increase of 5HIAA/5-HT ratio in the brain (Gesto et al., 2008; Gesto et al., 2013; Gesto et al., 2015; Øverli et al., 1999; Schjolden et al., 2006; Weber et al., 2012; Winberg and Nilsson, 1993). Serotonin plays a role in the HPI (and HPA) axis modulation and cortisol release, but its effects, activatory (Medeiros et al., 2010; Winberg et al., 1997) or inhibitory (Höglund et al., 2002; Saphier et al., 1995) remain unclear. Here, 10min after stress we detected a variation of brain 5-HT levels for mutant fish but not for wild type ones, both in SF and CF (Fig. Supp. 5B).

*Dopamine*: In fish, several studies demonstrated an activation of the dopaminergic system by a stressor (Gesto et al., 2008; Øverli et al., 1999; Weber et al., 2012), but this activation does not always take place and depends on the brain region (Weber et al., 2012), the social status (Øverli et al., 1999) and the stress duration (Gesto et al., 2015). It was proposed that DA activity could moderate the effects of the stress-induced serotonergic increase (Gesto et al., 2015; Höglund et al., 2001). Together, 5-HT and DA would be the first factors put in play by stressors and would regulate the intensity of the stress response (Gesto et al., 2015). In our study, brain DA concentrations did not vary after stress (Fig. Supp.

5C) -but we cannot conclude on an activation or not of the DA system since we did not access DOPAC levels.

A

| Location | Number of individuals | Number of polymophisms intra-location |
|---|---|---|
| Surface | 20 | 57 |
| Pachon | 20 | 23 |
| Los sabinos | 10 | 3 |
| Tinaja | 12 | 18 |
| Curva | 10 | 1 |
| Chica | 10 | 32 |
| Molino | 9 | 5 |
| Escondido | 7 | 1 |
| Caballo moro | 2 | 1 |
| Subterraneo | 10 | 13 |

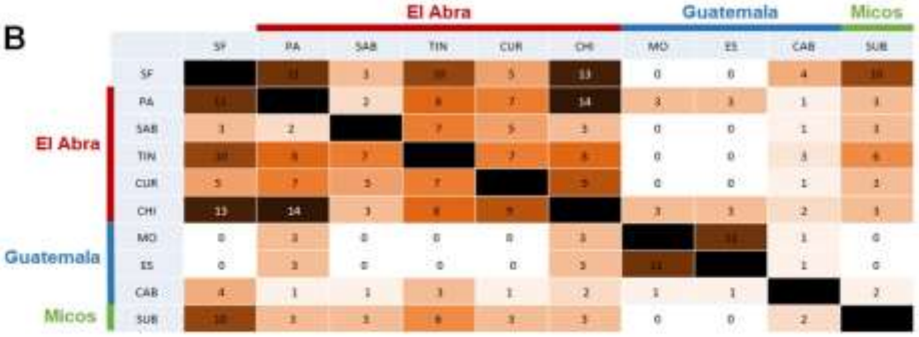


C

| | NC_035915. 1:32713783 | NC_035915. 1:32713768 | NC_035915. 1:32713768 | NC_035915. 1:32713733 | NC_035915. 1:32713703 | NC_035915. 1:32713347 | NC_035915. 1:32713238 | NC_035915. 1:32712805 | NC_035915. 1:32712378 | NC_035915. 1:32711901 | NC_035915. 1:32711590 |
|---|---|---|---|---|---|---|---|---|---|---|---|
| SF | TGACTAG | A | A | T | delA and WT | C and T | WT | G and T | C and T | C | A |
| Pa | AGCTA | T | C | T | delA | C | WT | T | C | C | A |
| Tin | TGACTAG | A | A and C | T and C | delA and WT | T | WT | G | T and C | T | A and C |
| Sab | TGACTAG | A | A | C | WT | T | WT | G | T | T | A and C |
| Cur | AGCTA | T | C | T | delA | C | deletion 7nt | G | T | T | |
| Chi | AGCTA | T | C | T | delA | C | deletion 7nt | T | C | T | A |

Supp 6

**Figure Supp 6. Polymorphism along a ~4 kb fragment of the *mao* gene, and *mao* haplotypes.**

(A) Number of wild-sampled individuals tested and number of polymorphisms in the ~4kb fragment around the P106L position in the *mao* gene, detected at each sampling site. For instance, we sequenced 20 fish from the river population, and we found 57 polymorphic positions distributed along the sequenced fragment. (B) Number of shared polymorphisms between surface populations and cave populations from El Abra, Guatemala and Micos groups. Darker colors indicate a high number of shared polymorphisms. Note that they are mostly shared within groups of caves. (C) Haplotypes reconstitution for the P106L mutant allele. Each column corresponds to a polymorphic position. The first line depicts the haplotypes encountered in SF (all Proline at position 106 – homozygote wildtype), and the next lines are haplotypes associated with the Leucine106 (P106L) mutant allele in cave populations of the El Abra group. For instance, all P106L mutant individuals sequenced from Pachón cave had the sequence AGCTA (first column in the table) at the position 32713783, whereas all P106L mutants from the Tinaja cave had the sequence TGACTAG instead at that same position. The P106L mutation is in position NC_035915.1:32711472.

**Video Supp 1: thigmotaxis in CF, 10min after being placed in a novel tank.**

**Video Supp 2: thigmotaxis in CF, 1hour after being placed in a novel tank.**

**Video Supp 3: thigmotaxis in CF, 96hours after being placed in a novel tank.**
